# Global Epistasis Constrains the Dynamics of Fitness in Multiple Environments

**DOI:** 10.64898/2026.11.14.766606

**Authors:** Jhelam N. Deshpande, Sarah M. Ardell, Sergey Kryazhimskiy

## Abstract

Adaptive mutations in one (“home”) environment often have diverse effects in other (“non-home”) environments. This so-called “pleiotropy” has important implications for the ability of populations to survive environmental fluctuations, which are common in nature. But how the fitness effects vary across mutations, genetic backgrounds and environments is poorly characterized empirically, and a unifying theory describing this variation is lacking. To address this gap, we develop the so-called “global epistasis” family of models and find that it explains a substantial proportion of variation in the distributions of mutational effects across genetic backgrounds and environments in a published yeast dataset. We then examine evolutionary dynamics that emerge in this model and we find that, as a population adapts at home, its non-home fitness can either converge to an intermediate finite value, or increase or decline indefinitely, with the empirical parameters being consistent with the first two outcomes. This work provides a quantitative unification of pleiotropy and epistasis, generates testable predictions for the pleiotropic consequences of adaptation and enables a better understanding of evolution in fluctuating environments.

## 1 Introduction

The ability of an organism to survive and reproduce—its fitness—fundamentally depends on the surrounding environment. Different environments place different demands on the organism and thereby introduce fitness trade-offs or trade-ups. Even in the absence of hard physiological constraints, positive or negative correlations between fitness in different environments can arise simply because certain phenotypes are more readily accessible by mutations. Regardless of the specific underlying mechanisms, we generally expect that a population adapting to one (“home”) environment would sustain some collateral fitness gains or losses in other (“non-home”) conditions. Such side effects of adaptation in one condition on fitness in another are an instance of pleiotropy, a term that broadly refers to situations when one mutation or gene affects multiple phenotypic traits (Paaby and Rockman, 2013; Wagner and Zhang, 2011). While collateral (or pleiotropic) fitness effects may not be “visible” to natural selection in the home environment, most organisms will eventually experience environmental shifts where the collateral effects of previously accumulated mutations would be exposed and may become crucial for the population’s long-term survival. Thus, understanding these effects is important for both fundamental and applied questions in ecology and evolution, such as maintenance of biodiversity (Kassen, 2002), mechanisms of ecological specialization (Futuyma and Moreno, 1988; Kassen, 2002), emergence of multidrug resistance (Andersson and Hughes, 2010; MacLean et al., 2010; Imamovic and Sommer, 2013; Pál et al., 2015; Hausser and Alon, 2020), persistence of populations in the face of climate change (Walworth et al., 2020; Willi and Buskirk, 2022; Salamin et al., 2010), the ability of viruses to infect novel hosts (Wang et al., 2024), etc.

Classical work in this area focused on fitness trade-offs, the idea that adaptation to one environment incurs fitness costs in others (Levins, 1962), often as an explanation for the diversity of species (Futuyma and Moreno, 1988) and local adaptation (Kawecki and Ebert, 2004). However, the more recent field and experimental studies paint a more complex picture (reviewed by Remold (2012); Bono et al. (2020); Visher and Boots (2020)). It appears that, although some genetic variants incur fitness trade-offs across environments (Hietpas et al., 2013), many others do not (Gifford et al., 2018), and still others incur collateral fitness benefits (Ostrowski et al., 2005; Jerison et al., 2020). Although the physiological causes underlying such variation in collateral effects of mutations are rarely known, some complexity can be expected because these effects must at least to some extent depend on the similarity between the home and non-home conditions (Jasmin and Zeyl, 2013; Leiby and Marx, 2014; Jerison et al., 2020; Bakerlee et al., 2021; Card et al., 2025) as well as on the proximity of the organism to the relevant Pareto front (Li et al., 2019).

While collateral fitness effects of mutations ultimately stem from how the organism’s physiology is encoded in its genome, some ability to predict and/or explain evolutionary outcomes may be achieved without such detailed mechanistic knowledge. In principle, to predict how fitness in home and non-home environments changes over time, it is sufficient to know the probability that a new mutation has certain effects on fitness in both conditions, a construct known as the joint distribution of fitness effects of mutations, or JDFE for short (Jerison et al., 2014; Ardell and Kryazhimskiy, 2021). Analogous to the one-dimensional distribution of fitness effects (DFEs), which dictates how fitness changes in one environment, the JDFE dictates changes in fitness in both home and non-home environments as the population adapts to the home condition. Specifically, the expected rate of fitness change at non-home during adaptation at home is captured by the so-called “pleiotropy statistic” of the JDFE, which is simply the effect of a typical mutation fixed at home on fitness in the non-home environment (Ardell and Kryazhimskiy, 2021). A positive value of the pleiotropy statistic indicates that the population’s non-home fitness is expected to increase during adaptation at home, while a negative value predicts the opposite.

Although the JDFE framework is general and potentially quite powerful, its practical utility hinges upon knowing the shape of JDFEs, or at least some of their properties, such as the pleiotropy statistic. However, not only do we currently lack estimates of empirical JDFEs, even for model organisms in lab conditions, there is currently little reason to believe that JDFEs should have one universal shape. Instead, there is overwhelming evidence that DFEs vary across environments, species and even genotypes of the same species (Martin and Lenormand, 2006; Eyre-Walker and Keightley, 2007; Qian et al., 2012; Blundell et al., 2019; Aggeli et al., 2021; Johnson and Desai, 2022; Ardell et al., 2024; Goldstein et al., 2025), indicating that the same must be true for JDFEs, as the former are the marginals of the latter. Since it is impossible to measure JDFEs across all relevant genotypes and conditions, we need a theory that would predict how JDFE shapes are expected to vary.

Over the past 15 years, many experimental studies, primarily in microbes, have indicated that DFEs vary across genotypes in a somewhat predictable manner. Specifically, more fit genotypes tend to have DFEs with fewer and weaker beneficial mutations (Barrick et al., 2010; Chou et al., 2011; Khan et al., 2011; Kryazhimskiy et al., 2014; Lukačišinová et al., 2020; Aggeli et al., 2021; Goldstein et al., 2025) and more abundant and stronger deleterious mutations (Johnson et al., 2019; Goldstein et al., 2025). This DFE variation appears to be driven at least in part by “global epistasis”, which refers to the fact that the fitness effects of individual mutations negatively correlate with the fitness of the genetic background into which they are introduced (Johnson et al., 2023). Furthermore, recent experimental work from our lab suggests that global epistasis can help explain variation in DFE shapes across environments as well (Ardell et al., 2024). This experimental work is consistent with a microscopic model proposed by Reddy and Desai (2021) and its multi-environment extension (Brooks et al., 2026).

Regardless of its mechanistic origins, since global epistasis helps us understand the variation among DFE shapes, it stands to reason that it may also help explain the variation among JDFEs, given that the former are the marginals of the latter. However, how global epistasis constrains JDFEs and how evolutionary dynamics unfold on such JDFEs is unknown. To address this gap here, we first use the data obtained by Ardell et al. (2024) to characterize the variation among empirical JDFEs across 30 ordered pairs of environments and 42 yeast strains. To explain this variation above and beyond what is known about the variation in DFEs, we extend the global epistasis model proposed by Reddy and Desai (2021) and Ardell et al. (2024) and obtain theoretical predictions for the JDFE covariance and the pleiotropy statistic. For any pair of environments, our statistical model describes a family of JDFEs that depends on 8 parameters. Finally, we explore the key features of evolutionary dynamics on this family of JDFEs, focusing in particular on how adaptation to a home environment affects non-home fitness.

## 2 Results

### 2.1 Empirical JDFEs vary across yeast strains and environments

We begin our investigation by examining how empirical JDFEs vary across genetic backgrounds and environments. To this end, we use our dataset described in Ref. (Ardell et al., 2024), which we refer to as the Ardell24 data. In that work, we measured the effects of about 100 quasi-random transposon-insertion mutations on growth rate in 42 strains of yeast *S. cerevisiae* in six environments that vary by pH (3.2, 5.0 and 7.0) and temperature (30^*°*^C and 37^*°*^C) in a factorial fashion.

A JDFE for a given genetic background in a given pair of environments is the probability distribution *J* (***s***) that a new mutation introduced into this genetic background has fitness effects (selection coefficients) ***s*** = (*s*_1_, *s*_2_)^*T*^ in these environments, with condition 1 usually being designated as “home” (Ardell and Kryazhimskiy, 2021). From the Ardell24 dataset, we constructed a total of 621 JDFEs across all strains and fifteen unique environment pairs in which there were at least 20 mutations. This filtering removes 9 strain-environment combinations. Figure 1A shows some representative empirical JDFEs for three strains and five pairs of environments from the Ardell24 dataset. This small but representative sample reveals that the shape of JDFEs varies substantially across both strains and environments.

**Figure 1.**
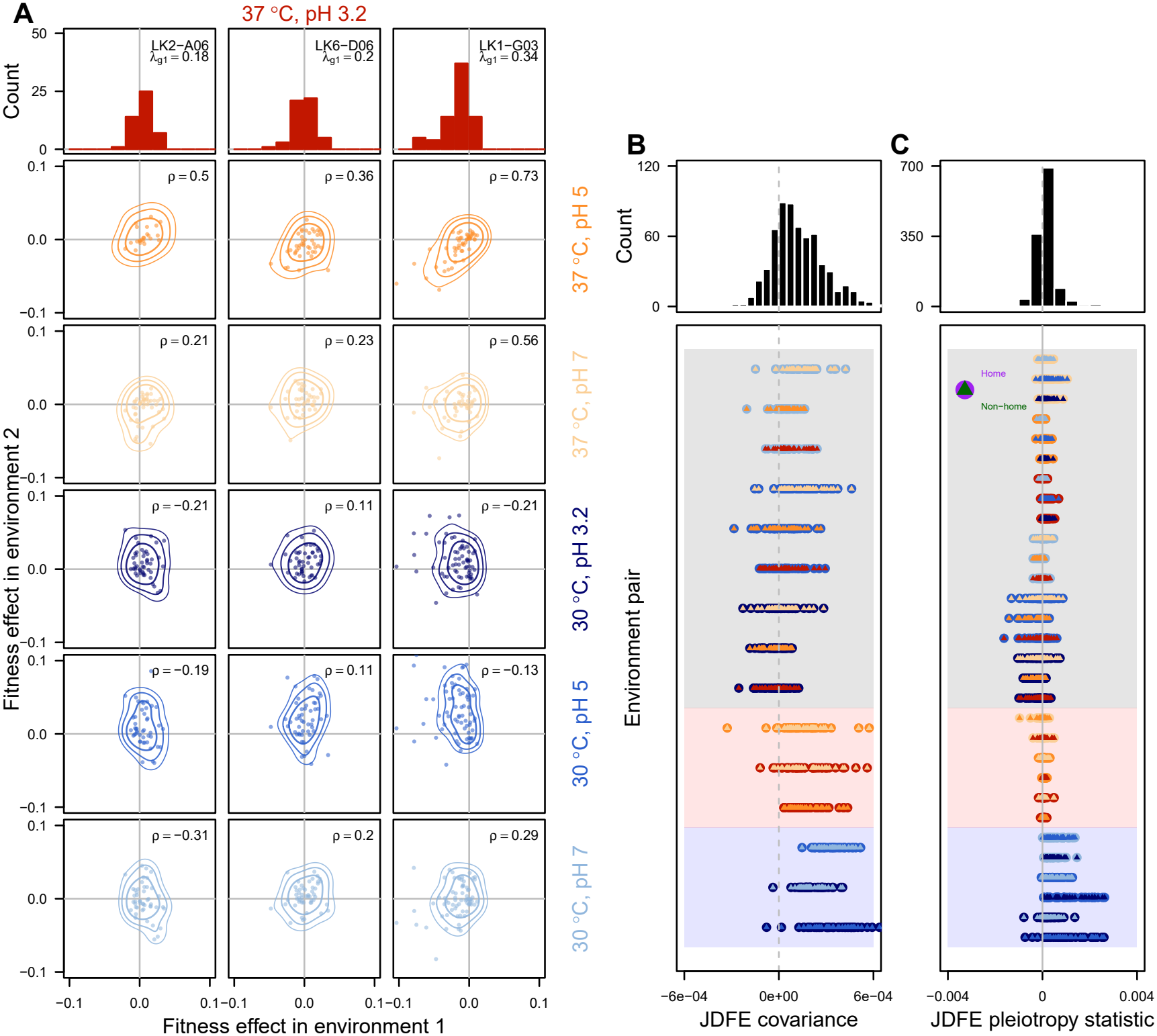
JDFEs vary across strains and environments in yeast. **A.** Top row shows the marginal distributions of fitness effects for three *S. cerevisiae* strains in environment 1, which is 37^*°*^*C* and pH 3.2. Each subsequent row shows empirical JDFEs with a different environment 2, as indicated. Each point represents a transposon-insertion mutation. Lines represent contours and the correlation coefficient is indicated on the top right. **B, C**. Histograms of replicate-averaged JDFE covariance and pleiotropy statistic with breakdown of variation in JDFE covariance and pleiotropy statistic by environment pair. The home environment is represented by the circle and non-home by triangle within the circle. Data are from Ref. (Ardell et al., 2024).

The Ardell24 dataset allows us to estimate a number of meaningful summary statistics of JDFEs, such as the mean vector, the covariance matrix, correlation coefficient, etc. The means and variances of the JDFE are properties of the marginal DFEs and have been previously characterized by Ardell et al. (2024). We therefore focus our analysis on properties that cannot be reduced to those of the marginal DFEs, namely JDFE covariance *c*^JDFE^ and the pleiotropy statistic mentioned in the Introduction, which we label *r*^JDFE^. Figure 1B,C show the distribution of JDFE *c*^JDFE^ and *r*^JDFE^. The mean JDFE covariance is positive (*⟨c*^JDFE^*⟩* = 1.2 × 10^*−*4^ and correlation *⟨ρ*^JDFE^*⟩* = 0.25). JDFEs in our dataset can be weakly negatively correlated or strongly positively correlated (95% percentile interval corresponding to [−1.2 × 10^*−*4^, 4.4 × 10^*−*4^] and [−0.21, 0.69] for JDFE correlation). The pleiotropy statistic (Figure 1C) in our dataset has mean ⟨*r*^JDFE^ ⟩= 1.4 × 10^*−*4^ and the range spans both negative and positive values (95% interval [−3.9 × 10^*−*4^, 8.9 × 10^*−*4^]).

We next used a linear mixed effects model to investigate how this variation is structured by the environment pair, strain and the interaction between them (Figure S1–S2; see Methods). We find that a substantial fraction of variation in JDFE covariance and pleiotropy statistic is attributable to environment pair, as seen in Figure 1B,C (38%, *p <* 0.001 for both). A smaller fraction is attributable to interaction between strain and environment (13%, *p <* 0.001 for *c*^JDFE^ and 22%, *p <* 0.001 for *r*^JDFE^). Residual between replicate variation accounted for 45% and 33% variation in *c*^JDFE^ and *r*^JDFE^, respectively. This breakdown of variance components is similar to the one we find for the properties of marginal DFEs (Figure S3–S4).

### 2.2 Generalized global epistasis explains the variation among empirical JDFEs

The fact that the interaction between the environment and the genetic background contributes substantially to the variation in JDFE properties could be interpreted as each strain having its own idiosyncratic JDFE in each pair of environments. However, Ardell et al. (2024) have shown that the fitness effects of individual mutations vary systematically with the growth rate of the genetic background. As a result, DFE moments vary predictably as a function of a strain’s growth rate. Thus, we asked if the same approach can be applied to explain the variation in JDFE covariance *c*^JDFE^ and pleiotropy statistic *r*^JDFE^.

In particular, Ardell et al. (2024) found that the fitness effect *s*_*mge*_ of mutation *m* that is introduced into genetic background strain *g* and measured in environment *e* can be statistically modeled as a linear function of the growth rate *λ*_*ge*_ of the background strain *g* in environment *e* as

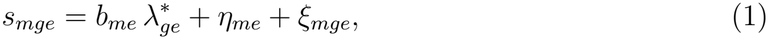

which is referred to as the generalized global epistasis model. Here, the predictor 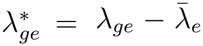 is the adjusted growth rate of the background strain *g* in environment *e*, i.e., its growth rate relative to the so-called “pivot” growth rate 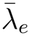 in environment *e* (discussed below), *b*_*me*_ is the global epistasis slope for mutation *m* in environment *e*, and terms *η*_*me*_ and *ξ*_*gme*_ represent the residuals, i.e., they account for variance in the fitness effects that cannot be explained by global epistasis. Specifically, the pivot growth rate 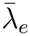 is interpreted as the growth rate of the background strain where a typical mutation switches sign in environment *e, η*_*me*_, referred to as the “pivot noise”, is a random variable that determines how far the global epistasis intercept of mutation *m* in environment *e* deviates from the average, and *ξ*_*mge*_, referred to as “idiosyncratic epistasis”, is a random variable that determines how much the effect of mutation *m* in the genetic background *g* in environment *e* deviates from the global epistasis trend. Ardell et al. (2024) found the estimated global epistasis slopes *b*_*me*_ to be overwhelmingly negative, such that their distribution had a negative mean and negative skewness. Moreover, they found these slopes to be largely invariant across environments, i.e., *b*_*me*_ ≈ *b*_*m*_ for all *e* (“same slope model”). For our theory, we consider a more general model that allows for these slopes to vary across environments (“different slopes model”). From here on, we assume that the growth rate is a proxy for fitness, and refer to 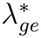 as adjusted fitness of strain *g* in environment *e*. We will discuss the limitations of this simplification later.

Ardell et al. (2024) considered the generalized global epistasis model (1) in each environment separately. To understand the implications of global epistasis on JDFEs, we need to consider equation (1) in two environments (*e* = 1 and *e* = 2) simultaneously. To this end, we assume that the global epistasis slopes ***b***_*m*_ = (*b*_*m*1_, *b*_*m*2_)^*T*^ are drawn from a bivariate distribution *p*_sl_ (***b***_*m*_), residuals ***η***_*m*_ = (*η*_*m*1_, *η*_*m*2_)^*T*^ and ***ξ***_*mg*_ = (*ξ*_*mg*1_, *ξ*_*mg*2_)^*T*^ are drawn from bivariate normal distributions with zero mean and covariance matrices **Σ**_pi_ and **Σ**_id_, respectively. We assume that ***b***_*m*_, ***η***_*m*_ and ***ξ***_*mg*_ are all mutually independent. Under these assumptions, we can express the JDFE *J*_*g*_ (***s***) for the background genotype *g* as a function of its fitness vector 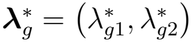 as

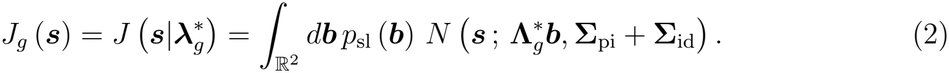

Here, *N* (***s***; ***µ*, Σ**) denotes a bivariate normal distribution with mean vector ***µ*** and covariance matrix **Σ**, and 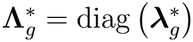 is the diagonal matrix generated by vector 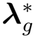.

Equation (2) allows us to compute various statistics of the JDFE as a function of the background fitness vector 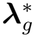 (see Section 4.3 in “Methods“). In particular, the covariance and the pleiotropy statistic of the JDFE in background *g* are given by

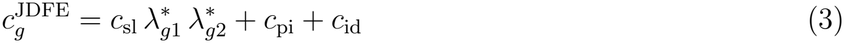

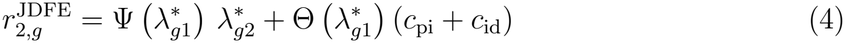

Here *c*_sl_ = Cov [*b*_*m*1_, *b*_*m*2_] is the cross-environment covariance of global epistasis slopes, *c*_pi_ = Cov [*η*_*m*1_, *η*_*m*2_] is the cross-environment covariance in the pivot noise, and functions 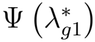 and 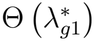 are given by equation (10) in “Methods” and visualized in Figure S5. Briefly, these two terms depend on the adjusted background fitness in the home environment, joint global epistasis slope distributions, the pivot and idiosyncratic noise in the home environment. Note that for the simplified model fit by Ardell et al. (2024), in which global epistasis slopes remain invariant, the covariance *c*_sl_ equals the variance in the slope distribution 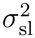.

Equations (3) and (4) show that, if mutations exhibit global epistasis (equation (1)), then the JDFE statistics should scale in a quantitatively predictable way with the adjusted fitness of the background strain. To test these quantitative predictions, we first estimate JDFE parameters using a linear mixed model (LMM) approach (see Section 4.2.2 in “Methods“). Equation (3) shows that, to understand the dependence of the JDFE covariance on fitness, we only need to estimate three JDFE parameters, *c*_sl_, *c*_pi_ and *c*_id_. For the pleiotropy statistic, we additionally need to know the distribution of global epistasis slopes, pivot noise and idiosyncratic variance in the home environment. Note that while our theoretical derivations assume arbitrary slope distribution, for inference, we assume that slopes follow a joint normal distribution across environments. We fit two models: a “different slopes” and “same slope” model, consistent with Ardell et al. (2024). Allowing slopes to vary between environments improved model performance at the level of predicting fitness effects, as expected because there are more parameters (SI Table S1). However, the different slopes model was not better at predicting JDFE covariance or pleiotropy statistic (SI Figures S6, S7 and S8). Therefore, in the main text, we only present results for the “same slopes” model. Our parameter estimates for both models are provided in SI Tables S2–S4 and Figures S6 and S9.

We found that the pivot noise terms and idiosyncratic epistasis terms are positively correlated across environments with the same temperature (correlation coefficient range [0.49,0.82] and [0.34, 0.78] for pivot noise and idiosyncratic terms, respectively) but nearly uncorrelated across environments with different temperatures (correlation coefficient range [0.04, 0.41] and [0.05,0.31] for pivot noise and idiosyncratic terms, respectively; Figure S9).

Substituting our parameter estimates into equations (3) and (4), we find that our predictions correctly capture the trends in how JDFE covariance and pleiotropy statistic change across strains and environments (Figure 2). In particular, given that 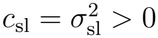 for all pairs of environments, we expect JDFE covariance to increase with the product of adjusted fitness 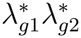, and this is indeed the case (Figure 2A). Furthermore, our model shows that the intercept of this relationship equals the sum of the cross-environmental covariance in pivot noise and idiosyncratic epistasis (see equation (3)). Given our estimates of *c*_pi_ and *c*_id_, we expect these intercepts to be positive in environmental pairs at the same temperature and nearly zero at different temperatures, and this is what we find (compare cold-versus warm-colored lines in each panel of Figure 2A). Overall, our model explains 49% of the variation in *c*^JDFE^, of which 31% is attributed to the environment pair (intercept) and 18.2% is explained by the global terms (Figure 2C).

**Figure 2.**
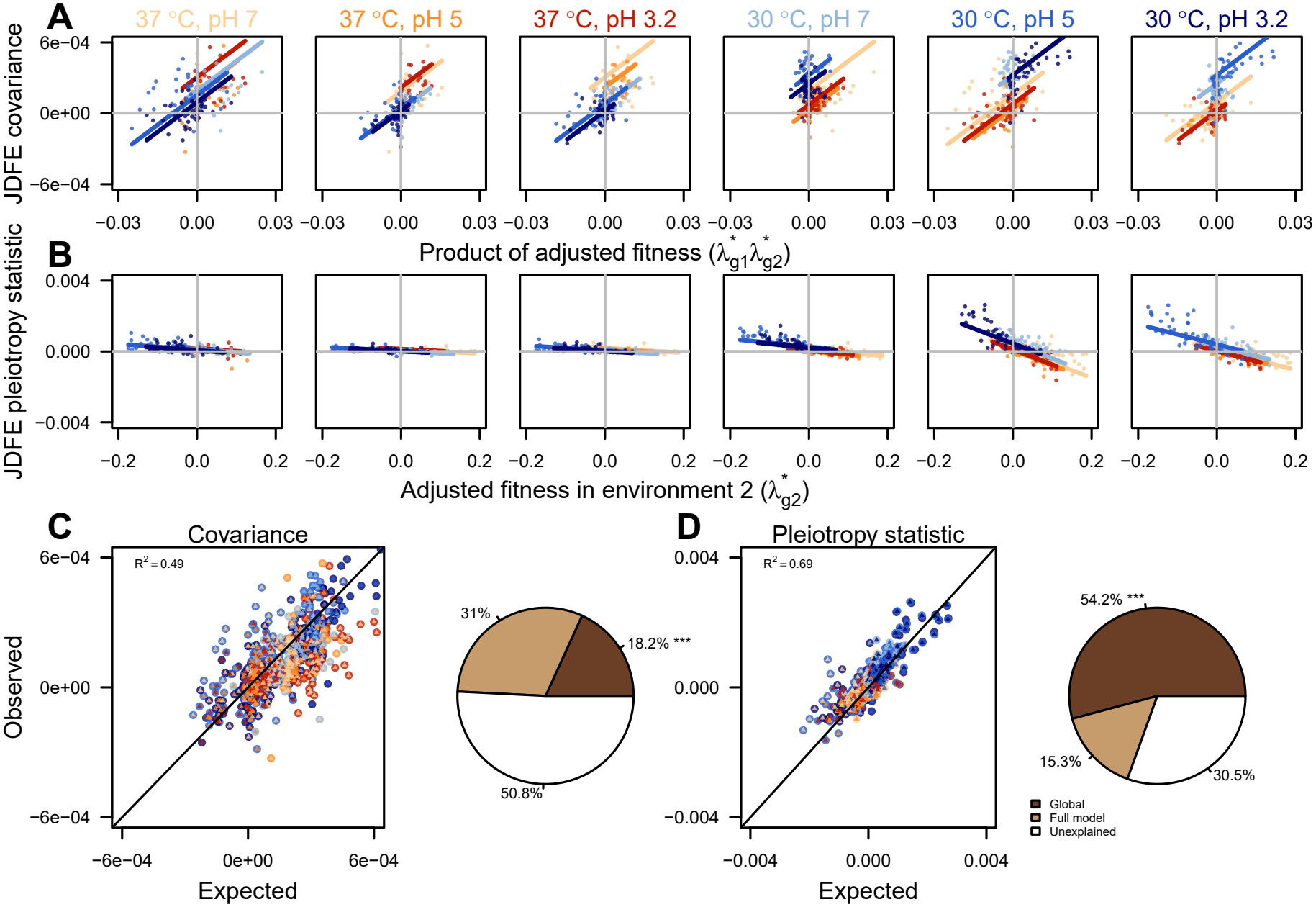
Global epistasis explains variation in JDFE statistics. **A.** JDFE covariance versus product of adjusted fitness. Each panel corresponds to one home environment. Each point represents a strain in a given pair of environments. Lines show predictions of the same slope global epistasis model. Colors (as indicated in the column labels) represent the non-home environment. **B**. Pleiotropy statistic versus adjusted fitness in non-home environment. Notations are the same as in panel A. Model predictions are evaluated at the median adjusted background fitness in the home environment for the corresponding environment pair. **C**. Observed versus expected JDFE covariance and incremental model performance including global and remaining terms. **D**. Observed versus expected JDFE pleiotropy statistic and incremental model performance including global and remaining terms.

We obtain similarly consistent results for the pleiotropy statistic. Our analytical calculations (see SI section S1.2.1 and Eq. (S28)) show that the slope Ψ should be a negative increasing function of 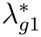 as long as the global epistasis slopes are negative (SI Figure S5). Therefore, based on our model (equation (4)), we expect the pleiotropy statistic *r*^JDFE^ to decrease with the non-home fitness 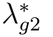 for any fixed value of 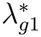. To visualize this relationship, we plot *r*^JDFE^ against 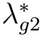 and find that the prediction line with slope 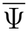 computed for the median value of 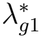 in the home environment (rather than an individual Ψ for each strain) already provides a good fit (Figure 2B). Furthermore, strains in this dataset tend to have higher median adjusted fitness at 37^*°*^C than at 30^*°*^C, which is expected to result in shallower scaling of *r*^JDFE^ with non-home fitness when the home environment is at 37^*°*^C than at 30^*°*^C. This is indeed what we observe (compare three left-most panels to the three right-most panels in Figure 2B). Quantitatively, our full model explains substantial variation in the pleiotropy statistic (*R*^2^ = 0.69; Figure 2D) of which the global-epistasis term 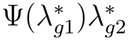 explains 54.2% and including the pleiotropic term 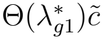 explains additional 15.3% variation.

Since our models (equations (3), (4)) depend on parameters that are estimated from the same data on which the model is tested, there is a potential for overfitting. We therefore carried out a cross-validation analysis in which we trained our model on 80% of strains and tested its predictions on the remaining 20% (see “Methods“, Section 4.4). We find that, in these out-of-sample predictions, equations (3) and (4) explain 43% and 69% of variation in *c*^JDFE^ and *r*^JDFE^, respectively (SI Figure S8). Thus, our generalized global epistasis model explains how JDFEs vary across strains and environments.

### 2.3 Properties of JDFEs induced by global epistasis

Equation (2) defines a family of JDFEs, which we refer to as the global-epistasis-induced JDFEs (or GE-JDFEs for short). This JDFE family is defined by the distribution of global epistasis slopes *p*_sl_ (***b***), and the covariance matrices **Σ**_pi_ and **Σ**_id_ for the pivot noise and id-iosyncratic epistasis. In this section, we explore some properties of GE-JDFEs. For simplicity, we focus in the main text on the case when both pivot noise and idiosyncratic terms are un-correlated across environments and slopes do not vary across environments. Characterization of other cases is provided in the Supplement (see SI Figures S10–S11).

We first examine the JDFE covariance. Equation (3) shows that JDFE covariance scales linearly with the product of adjusted fitness 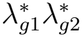, with the slope of this relationship being equal to the covariance of the slope distribution *c*_sl_. In the Ardell24 dataset, *c*_sl_ is positive for all pairs of environments (see SI Figure S6), which means that a genotype *g* has a positively correlated JDFE if its adjusted fitness has the same sign in both environments (Figure 3) and a negatively correlated JDFE if its adjusted fitness has the opposite signs in the two environments. This relationship has a simple intuitive explanation. First, the fact that global epistasis slopes are positively correlated implies that a mutation with a steep slope in one environment is likely to have a steep slope in the second environment. Now consider a background strain with both 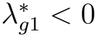 and 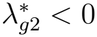 (symbol (−1, −1) in Figure 3). Since the global epistasis slopes are negative, mutations that are more beneficial in one environment tend to have steeper slopes and are therefore more likely to be beneficial in the second environment, which results in a positively correlated JDFE. The same logic applies to a background strain with both 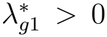 and 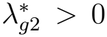 in which mutations with steeper slopes tend to be more deleterious in both environments (symbol (1, 1) in Figure 3). Now consider a background strain with 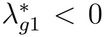 and 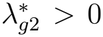 (symbol (−1, 1) in Figure 3). In this background strain, mutations with steeper slopes are more beneficial in environment 1 and more deleterious in environment 2, which results in a negatively correlated JDFE.

**Figure 3.**
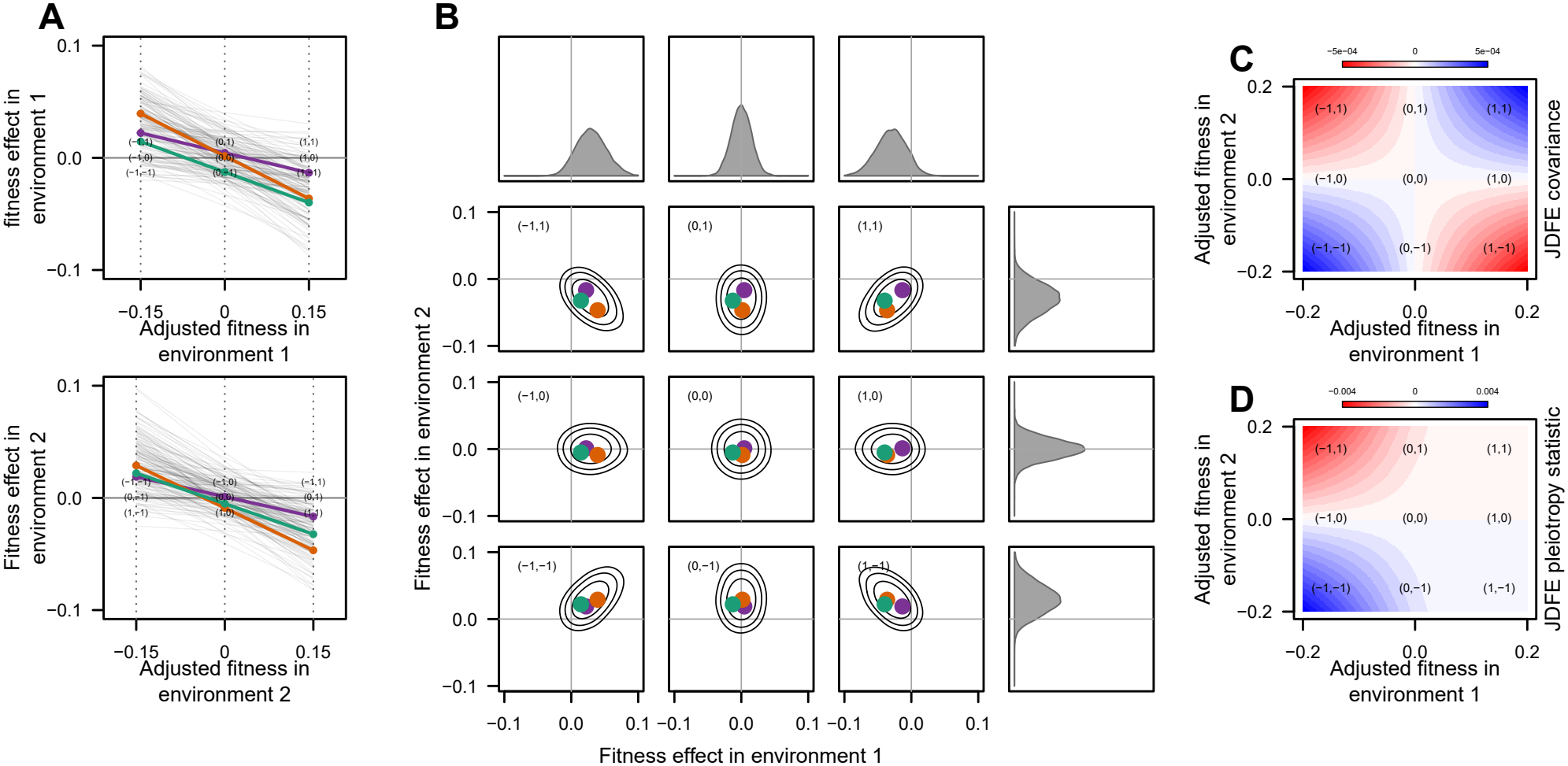
Schematic explaining how GE-JDFEs vary with home and non-home fitness in the absence of pleiotropic correlation. **A.** Mutations exhibit global epistasis with the same slope in both home (top) and non-home (bottom) environments. Three mutations are highlighted for illustration purposes. Dashed lines indicate three values of home and non-home fitness, below (−1), at (0), or above (+1) the pivot, which makes 9 combinations of background strains denoted by the combination of these labels. For example, (−1, −1) indicates a strain whose fitness is below the pivot in both environments. **B**. JDFE and both marginal DFEs for the 9 strains with different combinations of home and non-home fitness. Colored points correspond to mutations highlighted in panel A. **C**. Expected JDFE covariance across the plane of home and non-home fitness. **D**. Pleiotropy statistic across the plane of home and non-home fitness.

We now turn to the JDFE pleiotropy statistic. Equation (4) shows that *r*^JDFE^ scales linearly with the adjusted fitness in the non-home environment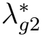 . The slope of this relationship Ψ depends on the adjusted background fitness in the home environment 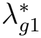 and on the distribution of global epistasis slopes *p*_sl_ (***b***). Loosely speaking, Ψ is negative when most global epistasis slopes are negative across the two environments, irrespective of the slope covariance *c*_sl_. Furthermore, it is an increasing function of the adjusted background fitness in the home environment 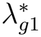. This relationship can be intuitively understood as follows. As increases, 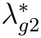 global epistasis makes all mutations on average less beneficial or more deleterious in the non-home environment, irrespective of their fitness effects in the home environment, which leads to a monotonically decreasing dependence of *r*^JDFE^ on 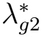 regardless of 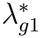, as long as the global epistasis slopes are negative. Additionally, the decreasing relationship of the absolute *r*^JDFE^ on the home fitness 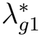 arises because, as 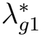 increases, adaptive mutations become less common and less beneficial at home, which reduces their fixation probability. Finally, we note that the intercept depends on both the pivot noise and idiosyncratic covariances (see Eq (4)), which will become important for understanding the pleiotropic consequences of long-term evolution in the home environment which we discuss in the next section.

In summary, this analysis reveals how the key features of the JDFE depend on the strain’s fitness in the home and non-home environments. In the absence of residual pleiotropic covariance, the sign of the JDFE correlation depends on whether a strain’s adjusted fitness in the two environments has the same sign. The pleiotropy statistic—which determines whether the population’s fitness in the non-home environment grows or declines as it adapts at home—depends primarily on the strain’s adjusted fitness in the non-home environment, such that genotypes with a positive adjusted non-home fitness have a negative pleiotropy statistic and vice versa.

### 2.4 Pleiotropic consequences of adaptation on JDFEs induced by global epistasis

As discussed above, the pleiotropy statistic *r*^JDFE^ determines how fitness in the non-home environment changes as the population adapts (Figure 4A) to the home environment (Ardell and Kryazhimskiy, 2021). Specifically, non-home fitness is expected to increase when *r*^JDFE^ *>* 0, and it is expected to decrease when *r*^JDFE^ *<* 0. Our analysis of equation (4) suggests that, on GE-JDFEs, these fitness dynamics follow qualitatively different patterns depending on the parameter 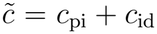, which captures the JDFE covariance not attributable to global epistasis (“pleiotropic covariance”).

**Figure 4.**
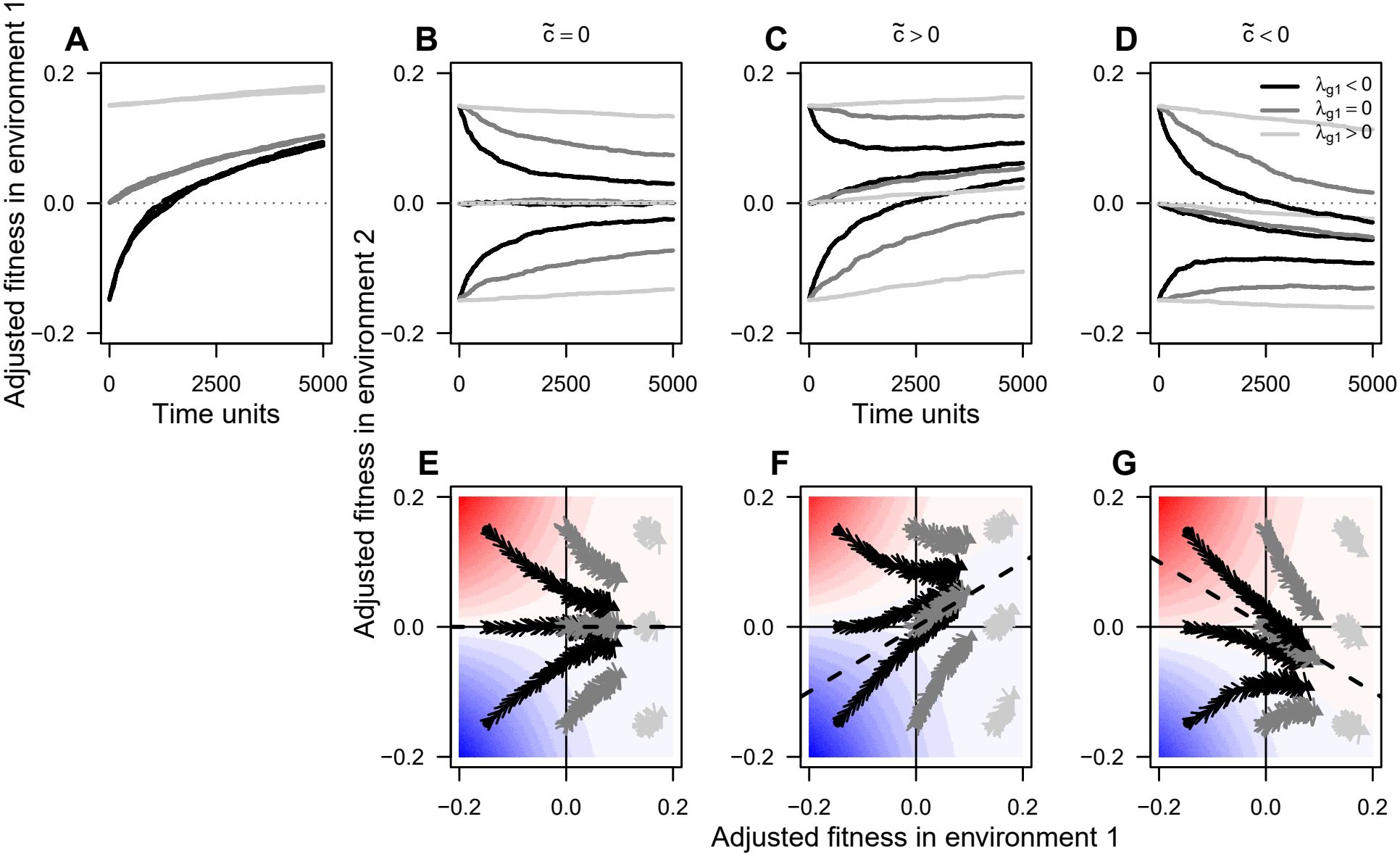
Evolutionary dynamics of fitness at the home and non-home environments on GE-JDFEs. **A.** Evolutionary trajectories of adjusted fitness in the home environment (environment 1), starting from three different initial fitness values 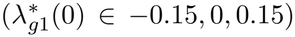. **B–D**. Corresponding trajectories of adjusted fitness in the non-home environment (environment 2) for zero (B), positive (C), and negative (D) residual pleiotropic covariance with nine initial conditions 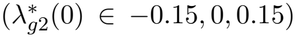. **E–G**. Evolutionary trajectories of adjusted fitness in environments 1 and 2, corresponding to panels A–D. Dashed lines indicate the attractor given by equation (5). Parameters: population size *N* = 10^4^, mutation rate *U* = 10^*−*5^, total number mutations that impact fitness pre-filtering for positive slopes *M* = 1000, mean of global epistasis slope distribution *µ*_sl1_ = *µ*_sl2_ = −0.16, standard deviation of GE slopes *σ*_sl1_ = *σ*_sl2_ = 0.13, correlation between GE slopes *ρ*_sl_ = 1, pivot noise standard deviation *σ*_pi1_ = *σ*_pi2_ = 0.02, idiosyncratic epistasis standard deviation *σ*_id1_ = *σ*_id2_ = 0.001, idiosyncratic epistasis correlation *ρ*_id_ = 0, and pivot noise correlation *ρ*_pi_ = 0, 0.5, −0.5 for panels B and E, C and F, and D and G, respectively. Global epistasis slopes are restricted to negative values.

We first consider the case 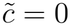, when both idiosyncratic and intercept terms in the home environment do not provide any information in the non-home environment. In the Ardell24 dataset, this situation is closest to environments at different temperature (Figure S9). In this case, equation (4) has a zero intercept, which implies that strains with zero non-home fitness have a JDFE with zero pleiotropy statistic. Furthermore, since generally Ψ *<* 0 in equation (4) (see SI Section S1.2.1 for a more nuanced discussion), the pleiotropy statistic and non-home fitness have the opposite sign. Therefore, the population’s non-home fitness is expected to decrease when it is positive, and it is expected to increase when it is negative, irrespective of the fitness in the home environment. Consequently, our theory predicts that, as a population adapts to the home environment, its fitness in the non-home environment is expected to converge to the pivot fitness 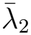 in that environment. We confirm this prediction using simulations of evolutionary dynamics in the strong selection weak mutation (SSWM) regime (see Section 4.5 in “Methods“). Interestingly, we find that the rate of convergence to the pivot fitness in the non-home environment strongly depends on the initial fitness at home.

Populations whose initial home fitness is below the pivot fitness at home adapt rapidly, which causes rapid concomitant changes in their non-home fitness. In contrast, populations that are already well adapted at home continue adapting slowly, which causes slow concomitant changes in their non-home fitness (compare dark and light gray arrows in Figure 4).

When 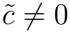, pivot fitness is no longer an attractor in the non-home environment. Below the home pivot, fitness trajectories are similar to the 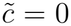 case (compare the left halves of panels E,F,G in Figure 4). However, once the population’s fitness crosses the home pivot, its non-home fitness begins diverging from the non-home pivot, either after crossing it or turning around. Equation (4) helps us understand these dynamics intuitively. It reveals that the fitness dynamics are governed by the balance of two “forces”. The first one is global epistasis, represented by the first term in equation (4): it pushes non-home fitness towards the pivot because it has the opposite sign to non-home fitness. The second “force” is the residual pleiotropic correlation, represented by the second term in equation (4), which drags the non-home fitness either along with the home fitness towards increasingly higher values (if 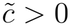, Figure 4C,F) or pulls it towards increasingly lower values (if 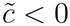, Figure 4D,G). The relative contributions of these two terms depend primarily on the fitness at home. When the home fitness is far below zero, the absolute value of Ψ is large, implying that global epistasis exerts a strong pull towards the non-home pivot. When home fitness is far above zero, Ψ ≈ 0, implying that global epistasis is weak and the dynamics are governed by the residual pleiotropic correlation.

It is possible to quantitatively determine the shape of the attractor to which the deterministic evolutionary trajectories are eventually expected to converge in the SSWM regime. In Section S1.2.2 of the SI we show that, if the global epistasis slopes of mutations are negative and invariant across environments, evolutionary trajectories converge to a line

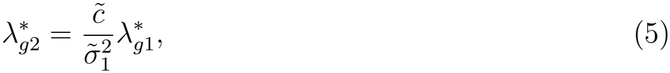

where 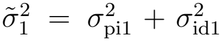 which is the total DFE variance in the home environment that is not attributable to global epistasis (“remaining variance”). Our simulations match this expectation (see Figure 4F,G).

Since the JDFEs in the Ardell24 dataset are well described by the invariant slope model, we estimated the slope of the attractor 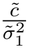 for all ordered environment pairs (Figure S14). We found that, on average, this slope was 0.64 for environments at the same temperature (range [0.25, 0.98]) and it was 0.17 for environmental pairs at different temperatures (range [0.05, 0.58]). This suggests that long-term adaptation of our yeast strains to one of our environments as its “home” would in general lead to concomitant pleiotropic fitness gains in all other environments from our set. These gains would be strong in environments where temperature is the same as at home, whereas fitness in environments with the other temperature would remain closer to the pivot.

## 3 Discussion

By constructing joint distributions of fitness effects (JDFE) from a large dataset of fitness effects of mutations in multiple genetic backgrounds across environments, we built a statistical model of cross-environmental global epistasis. It enabled us to understand how the two key JDFE properties, covariance and the pleiotropy statistic, depend on the fitness of the genetic background. These global epistasis relationships explain a substantial fraction of variation in the properties of empirically constructed JDFEs.

Going beyond the data presented here, the main consequence of reducing cross-environmental fitness effects of mutations to “global” or fitness-dependent components is that “diminishing returns” patterns that are observed within one environment generalize to cross-environmental fitness change. Specifically, as long as global epistasis constrains fitness effects of mutations to decrease with the fitness of a genetic background, and residual pleiotropic covariance is zero, saturating pleiotropic fitness increases and decreases are possible depending on the initial fitness of a genotype. Importantly, fitness in the non-home environment is expected to converge to the pivot fitness, which is a property of the non-home environment. When there is cross-environmental structure beyond global epistasis, as captured by positive or negative pleiotropic covariance, fitness trajectories are pushed away from the pivot.

Our work contributes to a broad understanding of cross-environmental pleiotropic evolution by identifying two major constraints to a generalization of “diminishing returns” adaptation to cross-environmental evolution: First, a genotype-to-fitness mapping that across distinct environmental conditions preserves negative global epistasis, or the tendency of mutational fitness effects to covary negatively with background fitness. Second, the residual pleiotropic covariance, which captures how distinct environments modulate fitness effects of mutations beyond global epistasis. Thus, for our results to be applicable to empirical systems, they should satisfy these constraints. Below, we discuss our main results in light of both theoretical and empirical evidence for preservation (or not) of negative global epistasis across environments and identifying the role of pleiotropic environment-specific effects. Finally, we synthesize the implications of our findings for cross-environmental fitness change and discuss some empirical work that could provide support for our predictions.

### 3.1 Preservation of global epistasis across environments is critical for the predictability of pleiotropic fitness change

In our study, we took a statistical approach to modeling cross-environmental global epistasis without positing a mechanism. Thus, potentially different kinds of genotype-to-fitness maps can explain our data since global epistasis with primarily negative slopes can emerge in several ways reviewed by Diaz-Colunga et al. (2023b). However, the residual structure of the data presented here (Ardell et al., 2024) aligns with work (Lyons et al., 2020; Reddy and Desai, 2021; Cotto and Day, 2023; Brooks et al., 2026) that suggests that at least within an environment, global epistasis is likely to emerge as “regression towards the mean” for genotype-to-fitness maps with certain features (Diaz-Colunga et al., 2023b), for instance those in which fitness of close genetic backgrounds are relatively uncorrelated (Reddy and Desai, 2021). Recently, this theory has been generalized to model cross-environmental global epistasis and evolution in fluctuating environments and could, in principle, be used to derive equivalent results. In particular, the widespread random epistasis (WRE) regime provides a possible microscopic basis for the preservation of negative global epistasis across environments, although the resulting pleiotropic structure also depends on how genetic interactions are shared across environments (Brooks et al., 2026).

Thus, if “regression towards the mean” indeed explains widespread “diminishing returns” adaptation, it is quite generic and is likely to be preserved across environments. As a consequence, we predict that our generalization of classically observed “diminishing returns” adaptation to cross-environmental fitness change should hold for the systems in which such adaptive trajectories have been repeatedly observed across multiple environments.

Limits to our theory may arise if environments break the tendency of fitness effects of mutations to covary negatively with fitness. For instance, Diaz-Colunga et al. (2023a) found for their set of four mutations underlying anti-malaria drug resistance in *Plasmodium falciparum* that global epistasis slopes go from being negative to positive for some mutations depending on drug concentrations. Similarly, recent theoretical work (Venkataraman and Saini, 2025) also indicates that metabolic models of global epistasis may not fall into a “regression towards the mean” framework and rather might be dependent on the frequency of mutations in certain pathways. Thus, if these pathways are more or less important depending on environments, the tendency for global epistasis slopes to be primarily negative could break. Additionally, such mechanistic models could produce additional structure in residuals (Reddy and Desai, 2021; Brooks et al., 2026) not captured by our model. Such empirical and theoretical contradictions clarify that our theory should be applicable to environments that do not fundamentally change the structures of the genotype-to-fitness map producing global epistasis, which at least appears to be the case for data from Ardell et al. (2024) presented here.

### 3.2 Residual pleiotropic correlation as a measure of environmental similarity

Although we considered only six environments, our analysis indicates that the residual pleiotropic covariance between mutational fitness effects beyond global epistasis provides a potential measure of environmental similarity. In our data, this covariance was strongly positive between environments at the same temperature but weaker between environments at different temperatures, suggesting that mutations with unusually beneficial effects in one environment tend to have similarly beneficial effects in more similar environments. This residual covariance becomes particularly important later in adaptation, when the contribution of global epistasis to pleiotropic fitness change is reduced. More generally, our results suggest that collateral fitness gains or losses need not arise from strict physiological trade-offs or trade-ups between environments, but are instead a result of a shared fitness dependence of mutational effects. Thus, the direction of pleiotropic fitness change depends both on the fitness of the genetic background and on the extent to which mutational effects are shared across environments.

It must be mentioned here that our predictions differ from those of the Fisher’s geometric model (FGM; Tenaillon 2014) which has guided past empirical work on cross-environmental fitness change during adaptation (Jasmin and Zeyl, 2013; Olazcuaga et al., 2021). The FGM has also been proposed as an explanation for declining adaptability (Tenaillon, 2014) since fitness effects become smaller closer to the optimum. Indeed, Martin and Lenormand (2015) analyzed the FGM with universal pleiotropy for cross-environmental fitness change: here, background fitness dependence in selective and non-selective environments also occurs as a consequence of distance to an optimum corresponding to those environments. Thus, pleiotropic fitness losses are inevitable even though transient increases are possible. Interestingly, our model could capture similar qualitative behavior if we have negatively correlated pleiotropic residuals.

### 3.3 Comparison with empirical studies of cross-environmental evolution

While our statistical model explained cross-environmental properties of global epistasis-constrained JDFEs, particularly the scaling of the pleiotropy statistic 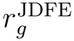 with background fitness, a definitive test of our theory remains. This is because it is possible that transposon insertion yeast mutants are in some way special or biased towards producing cross-environmental global epistasis. Thus, empirical validation requires experimental evolution in one environment and cross-environmental fitness assays. In the absence of this empirical test, we can compare some existing studies with the required experimental design to our predictions. Recent work by Ahlawat et al. (2025) in *E. coli* also indicated that starting fitness in the non-selective environment was a predictor of pleiotropic fitness change for a set of non-synonymous sugar environments, and consistent with our predictions, fitness gains were larger when ancestral fitness was lower in the non-home environment.

The study by Bakerlee et al. (2021), also on *S. cerevisiae*, reported diverse cross-environmental pleiotropic trajectories in distinct environments. We cannot directly test our predictions on this dataset since they measure competitive fitness relative to an ancestor whereas we used data for absolute growth rates. However, we can compare qualitative dynamics of fitness to that of our model. Broadly, they observed rapid initial fitness gains followed by declining adaptability in home environments, while fitness trajectories in non-home environments included monotonic gains or losses and non-monotonic changes, often involving initial fitness gains followed by subsequent declines. Our model captures these qualitative classes of trajectories by varying initial fitness in the non-home environment and the pleiotropic covariance 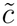 (see SI Fig. S15). Therefore, our model has some support from empirical studies but a definitive test remains.

### 3.4 Conclusion

Taken together, our work derives predictions that generalize typical “diminishing returns” adaptation in home environments to identify corresponding distinct classes of long-term trajectories of pleiotropic evolution in non-home environments. These evolutionary outcomes arise from classes of GE-JDFEs that we find in this study explain the variation in a large empirical data set of transposon insertion mutations (Ardell et al., 2024). Pleiotropic trajectories depend qualitatively on the initial fitness in the non-home environment and the residual pleiotropic correlation beyond global epistasis. As such, our model should be considered a baseline expectation and deviations from predicted trajectories could potentially represent additional structure not captured by fitness alone. More broadly, our work provides a statistical alternative to strict physiological trade-offs as an explanation for collateral fitness decreases and increases.

## 4 Methods

### 4.1 Data

Yeast mutant data were obtained from Ref. (Ardell et al., 2024) where a detailed description of how the data were produced can be found. Briefly, Ardell et al. (2024) used 42 strains of yeast *Saccharomyces cerevisiae* originally created by Bloom et al. (2013) as “background” strains for subsequent mutagenesis. These strains are derived from a cross between the lab strain BY and the wine strain RM and vary at approximately 40 thousand loci (Bloom et al., 2013; Johnson et al., 2019). Approximately 100 barcoded transposon insertion mutations (tn-mutations for short) were introduced into each background strain. Growth rates of the resulting mutants were measured in 6 conditions with two-fold biological replication. All conditions were based on the synthetic complete (SC) medium and varied by temperatures (30^*°*^C and 37^*°*^C) and pH (3.2, 5 and 7).

From this dataset, we obtained post-filtering estimates of the background growth rates *λ*_*gre*_ and the fitness effects of mutations *s*_*mgre*_ which are denoted by Δ*λ*_*mgre*_ in Ref. (Ardell et al., 2024). Indices *m, g, r* and *e* refer to the mutation, background strain, replicate and environment, respectively.

### 4.2 Data analysis

#### 4.2.1 Statistics of empirical JDFEs

The Ardell24 dataset allows us to construct empirical JDFEs for 42 yeast strains in 30 ordered pairs of environments. The total number of ordered empirical JDFEs is 1242 (621 unordered), which is less than 42 × 30 = 1260 because some strains dropped out of some environments. These empirical JDFEs contain between 37 and 83 mutations (median 72).

Thus, for each genetic background *g* and a pair of environments *e*_1_ and *e*_2_, we obtain the estimate of the covariance *c*^JDFE^ and the pleiotropy statistic 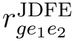. We estimate the pleiotropy statistic as 2*P* [*s*_1_ > 0] *⟨s*_1_*s*_2_|*s*_1_ > 0⟩ where ⟨·⟩ denotes the mean over all mutations (Ardell and Kryazhimskiy, 2021). We obtain 621 independent estimates of JDFE covariance (since these are symmetric with respect to environments) and 1242 estimates of the pleiotropy statistic. To understand how the variation was structured for a given JDFE parameter, we fit a linear mixed-effects model in R using lme4, of the form *x* ∼ environment pair+(1|strain) + (1|environment pair : strain). Environment pair was treated as fixed, strain and environment-pair-by-strain effects as random, and fixed-effect significance was assessed using lmerTest.

#### 4.2.2 Inference of global epistasis parameters from the Ardell24 dataset

In our original study (Ardell et al., 2024), we analysed the transposon-insertion mutagenesis data in yeast using a series of ordinary least squares (OLS) linear regressions. We found that the inferred global epistasis slopes were overwhelmingly negative, nearly invariant across environments and slopes correlated negatively with intercepts in each environment. These observations led us to propose the generalized global epistasis model (equation (1) with *b*_*me*_ = *b*_*m*_) as the most parsimonious explanation for the data.

However, the OLS approach does not directly estimate cross-environment correlations for the global epistasis slopes, pivot noise terms and idiosyncratic epistasis terms. Furthermore, it is subject to two statistical artifacts. First, as pointed out by Berger and Postma (2014), there is a negative bias in the correlation between the estimated fitness effects of mutations and the estimated background fitness. This bias arises when the fitness effects of a mutation in different backgrounds are estimated as differences between the growth rates of mutants and the growth rates of background strains, in which case the background growth rate appears on both sides of the linear regression equation. Second, OLS estimators of slope and intercept are in general themselves correlated (Seber and Lee, 2003). Although our previously reported results were robust with respect to several checks, it is desirable to develop an approach that would not only be more statistically sound but would also allow us to estimate the aforementioned cross-environment correlations.

Here, we describe such an approach within the linear mixed model (LMM) framework, and use it to reanalyze the Ardell24 dataset. First, we outline the logic of our approach in the idealized case where the true values of the fitness effects of mutations *s*_*mge*_ and the background growth rates *λ*_*ge*_ are known, where indices *m, g* and *e* denote the mutation, genetic background and environment, respectively, as before. Note that this model is not actually fit to the data but rather is used to motivate the analysis. We then extend this model to the realistic case of fitness effects and background growth rates measured with errors.

##### Model without measurement errors

In this idealized case, we could infer the global epistasis parameters by fitting the LMM

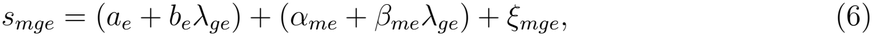

where the two parenthesis terms represent fixed and random effects, respectively. In this model, it would be reasonable to assume that the random slopes and intercepts are independent for different mutations but can covary for the same mutation. This model can be written in the Wilkinson notation (Wilkinson and Rogers, 1973) as *S ∼* 0+*e*+Λ : *e*+(0+*e*+Λ : *e*|*m*). This model would estimate the covariances between slopes and intercepts across mutations 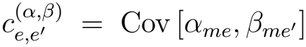 in each pair of environments *e, e*^*′*^. The statistical significance of these covariances could be tested using the likelihood ratio test against the nested model in which we set 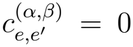, which could be written in the Wilkinson notation as: *S ∼* 0 + *e* + Λ : *e* + (0 + *e*|*m*) + (0 + Λ : *e*|*m*). If 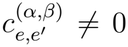, then *α*_*me*_ can be expressed as 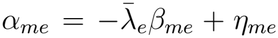 where 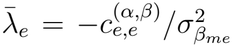 is the background growth rate at which the global epistasis regression lines for different mutations intersect each other, referred to as the “pivot” background growth rate (Ardell et al., 2024), 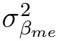 is the variance in *β*_*me*_ in environment *e*, and *η*_*me*_ are random variables with mean zero that are uncorrelated with *β*_*me*_, referred to as the “pivot noise”. Furthermore, if 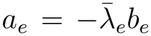, then the pivot point lies on the abscissa, i.e., an average mutation switches sign at 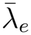. In this case, model (6) simplifies to equation (1) where *b*_*me*_ can be treated as either fixed or random effects. Allowing global epistasis slopes to vary among mutations and environments as random effects, model (1) can be written in Wilkinson notation as *S ∼* 0 +Λ^*\**^ : *e* +(0 +Λ^*\**^ : *e*|*m*) +(0 + *e*|*m*) +(0 + *e*|*m* : *g*).

Here, (0+Λ^*\**^ : *e*|*m*) describes the covariance of mutation-specific global epistasis slopes across environments, (0 + *e*|*m*) describes the covariance matrix **Σ**_pi_ of pivot noise, and (0 + *e*|*m* : *g*) describes the covariance matrix **Σ**_id_ of idiosyncratic epistasis. Below, we extend this model to account explicitly for measurement error in the estimated growth rates.

##### Model with measurement noise

Since the true values of fitness effects of mutations and background growth rates are unknown, and we only have access to their estimated values, model (6) is insufficient. To address this problem, denote the true growth rate of mutant *m* by *λ*_*mge*_ and its value estimated in replicate *r* by 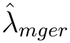. We assume that 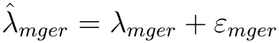 where *ε*_*mger*_ are the measurement noise terms, which we assume to be independent and identically distributed (i.i.d.) random variables with zero mean and environment-specific variance 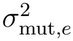. Similarly, the background growth rates estimated in replicate *r* are 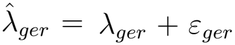 with *ε*_*ger*_ being i.i.d. with zero mean and environment-specific variance 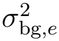 The fitness effect of mutation *m* is defined as *s*_*mge*_ = *λ*_*mge*_ − *λ*_*ge*_ and its estimated value is *ŝ*_*mger*_ = *s*_*mge*_ + *ε*_*mger*_ − *ε*_*ger*_. Using equation (6), we obtain the model for *ŝ*_*mger*_,

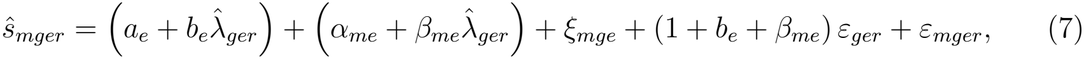

where we replaced −*ε*_*ger*_ with *ε*_*ger*_ assuming that they come from a distribution symmetric around zero. Model (7) does not fit into the standard LMM framework because of the product of fixed and random-effect parameters in the fourth term. Therefore, instead of fitting this model directly, we construct a series of simpler standard LMMs.

First, we estimate the correlation between global epistasis slopes and intercepts in each environment separately using the model

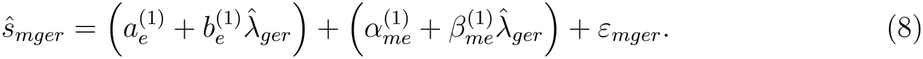

This model can be written in the Wilkinson notation as *S ∼* 1 + Λ + (1 + Λ|*m*). This model estimates the covariance 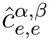 between 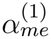 and 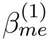 for each environment *e*. We test the statistical significance of these correlations against the nested model *S ∼* 1 + Λ + (1|*m*) + (0 + Λ|*m*) using the likelihood ratio test. Since the model (8) with correlated slopes and intercepts has six parameters in each environment and the nested model without correlations has five parameters, the relevant *χ*^2^ distribution has 1 degree of freedom. If the slope-intercept correlation in environment *e* is statistically significant, then we estimate the pivot growth rate as 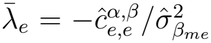, where 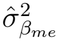 is the variance of 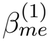 in environment *e*. Using these pivot growth rates, we compute the adjusted growth rates for all background strains as 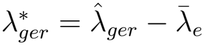. Then we use the second model

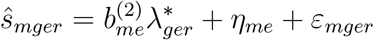

to obtain the initial estimates of the global epistasis slopes. This model can be written in the Wilkinson notation as *S ∼* 0 + Λ^*\**^ : *m* : *e* + (1|*m* : *e*). It returns the environment-specific estimates of slopes 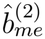 for each mutation.

Finally, we use these slope estimates to construct the random-effects design matrix in our final model,

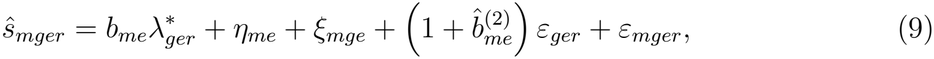

which we can write in the Wilkinson notation as *S ∼* 0 + Λ^*\**^ : *m* : *e* + (0 + *e*|*m*) + (0 + *e*|*m* : *g*)+(0+*B*2 : *e*||*g* : *r*) where the last term *B*2 is a vector of predictors 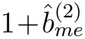 and “||” indicates that the slopes *ε*_*ger*_ on this predictor are independent and distributed with environment-specific variances 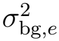. This model estimates the mutation and environment-specific slopes *b*_*me*_, the pivot noise covariance matrix **Σ**_pi_, the idiosyncratic epistasis covariance matrix **Σ**_id_, and the vectors of environment-specific error variances 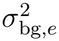 and 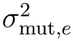.

##### Implementation

We implemented our approach using the `R` package `glmmTMB` for two models, one assuming global epistasis slopes remain invariant across environments and another in which they can vary. The full code will be made available in the deposited data, the SI Methods contains a summary of the `glmmTMB` formulas.

### 4.3 Deriving GE-JDFEs and their statistics

Starting from equation (1) in the main text, for a given mutation *m*, ***s*** = (*s*_*mg*1_, *s*_*mg*2_) denotes the joint fitness effect of that mutation in environments 1 and 2 for a strain *g*. Generalizing Ardell et al. (2024)’s approach to environment pairs, we assume that the pivot noise and idiosyncratic residuals ***η*** = (*η*_*m*1_, *η*_*m*2_) and ***ξ*** = (*ξ*_*mg*1_, *ξ*_*mg*2_) are drawn from joint normal distributions with zero means and a variance-covariance matrix **Σ**_pi_ (variance 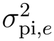 for environment *e* and *c*_pi_ for a pair of environments) and **Σ**_id_ (variance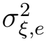 for environment *e* and *c*_id_ for a pair of environments). For the general theory, we assume that a cross-environmental pair of global epistasis slopes ***b*** = (*b*_*m*1_, *b*_*m*2_) is drawn from an arbitrary bivariate distribution *p*_*sl*_(***b***). Thus, for a single mutation *m* in a strain *g*, the joint fitness effect is drawn from a bivariate normal with mean vector 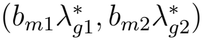 and a variance-covariance matrix **Σ**_pi_ +**Σ**_id_. So for a collection of mutations, the JDFE is given by Eq (2). From this distribution, we can calculate any property 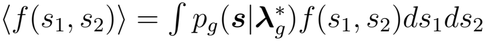. Thus, covariance is calculated as *c*_*g*_ = *⟨*(*s*_1_ −*⟨s*_1_*⟩*)(*s*_2_ −*⟨s*_2_*⟩*)*⟩* and the pleiotropy statistic as 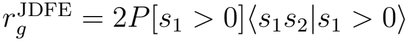 to obtain equations (3) and (4) with 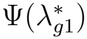 and 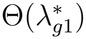 defined as:

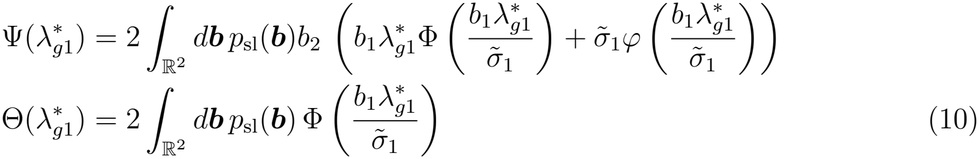

with *φ*(*z*) denoting the standard normal density and Φ(*z*) the standard normal cdf. See SI methods for detailed derivation and analysis.

### 4.4 Evaluating model performance

We used parameter estimates from the modified LMM described in 4.2.2 and substituted them in equations (3) and (4). The terms in Eq. (10) were calculated by Monte Carlo integration. We then used a linear model to estimate *R*^2^ between observed and model-predicted quantities using the `lm` function in `R` (version 4.1.2). All analyses were performed for the same slope and different slope models.

#### 4.4.1 Partitioning global vs. residual pleiotropic components

We fit sequential linear models and calculated corresponding *R*^2^ to understand the relative contribution of the terms of the model. We first fit a model using the global terms containing non-home fitness in equations (3) and (4). The 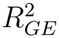 for this model represents the contribution of the global term and the 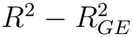 the variation explained by the remaining terms. We note that this depends on the order in which these terms are included, and that this is not a variance partition since the terms are correlated.

#### 4.4.2 Cross-validation analysis

To assess the predictive performance of our global epistasis model on genetic backgrounds not used for parameter estimation, we performed a strain-level holdout analysis. We randomly divided the 42 yeast strains into a training set of 34 strains (80%) and a test set of 8 strains (20%). We fit the full model described in 4.2.2, including estimation of the pivot using the training strains and evaluated predictions on the held out strains.

### 4.5 Evolutionary Simulations

#### 4.5.1 JDFE

We assume that there are a total of *M* = 1000 mutations that can potentially impact fitness across the two environments. Each mutation is related to the adjusted fitness in environments 1 and 2 through the generalized GE equation (1). We assume that the slopes *b*_*me*_ are drawn from a univariate or multivariate normal distribution, truncated to negative values, depending on whether we are simulating the same slope or variable slope scenarios. Similarly, the pivot noise terms are drawn from a multivariate normal with mean 0, and correlation coefficients corresponding to the three scenarios in the main text (see Figure 4 for parameters). Since idiosyncratic epistasis could in principle depend on the genotype, we redraw these terms each time step.

#### 4.5.2 SSWM simulations

We simulate evolution in the Strong Selection Weak Mutation (Gillespie, 1983) regime using the Gillespie algorithm (Gillespie, 1977). Briefly, we sample the waiting time until the next mutation from an exponential distribution with rate *NU*, where *N* is the population size and *U* is the mutation rate. In our simulations, *NU* = 0.1. We randomly sample 1 mutation with fitness in the home and non-home from this JDFE. This mutation then has a fixation probability of:

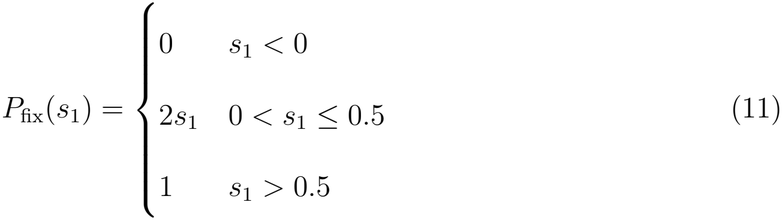

If the mutation fixes, we update fitness in the home and non-home. We continue the cycle until the population reaches the desired total number of generations.

## S1 Supplementary methods

### S1.1 Deriving the joint distribution of fitness effects (JDFE) from the global epistasis equation

Our objective is to understand how fitness of a population changes in an arbitrary environment *e* = 2 during adaptation to another environment *e* = 1 when there is global epistasis in the fitness effects of mutations across environments. To predict such trajectories (under assumption of strong selection weak mutation; SSWM) we need to know: a) the current fitness of the (clonal) population composed of a strain *g* in environments *e* = 1 and *e* = 2 (*λ*_*g*1_ and *λ*_*g*2_) and b) the joint distribution of fitness effects (JDFE) *P*_*g*_[*s*_1_, *s*_2_] which will determine how much fitness changes in both environments. The fitness effect of a mutation *s* for a given environment is defined as the difference between the mutant and wildtype growth rate. Since there is global epistasis, *P*_*g*_[*s*_1_, *s*_2_] also depends on *λ*_*g*1_ and *λ*_*g*2_, we should be able to predict pleiotropic trajectories with information only about initial growth rates in both environments 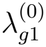 and 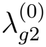 and statistical features (slopes, intercepts, etc.) of global epistasis. Thus, we will use the results of Ardell et al. (2024) as a starting point to build a generalized model for global epistasis in fitness effects of mutations across environments and derive a background growth rate dependent joint distribution of fitness effects (JDFE).

We write, for a given mutation *m* in strain *g* and environment *e*, the fitness effect *s*_*mge*_ as a function of 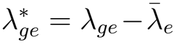, which is the background growth rate of the strain adjusted by the pivot growth rate for a given environment. By definition of pivot growth rate, 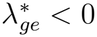 implies that mutations in strain *g* and environment *e* are primarily beneficial and 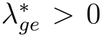 implies that they are primarily deleterious. The generalized global epistasis equation is written as:

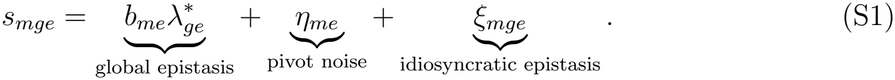

Here *b*_*me*_ is the slope with which the fitness effect of a mutation decreases with the adjusted background fitness for a given environment *e* representing the “global” component of epistasis, *η*_*me*_ is an environment specific intercept and *ξ*_*mge*_ is an idiosyncratic strain, environment and mutation specific residual effect. Note that this equation differs from Ardell et al. (2024) since we allow for the slope *b*_*me*_ to depend on both the mutation and environment, rather than just the mutation. Ardell et al. (2024) found that a common slope can explain the data well for global epistasis within an environment. Yet, we opt for the more general model to build our theory which allows us to a) extend our theory beyond the specific features of our dataset and b) test whether the explanatory power of a single slope model also extends to a cross-environmental analysis.

Denote ***s***_*mg*_ = (*s*_*mg*1_, *s*_*mg*2_)^*T*^, ***b***_*m*_ = (*b*_*m*1_, *b*_*m*2_)^*T*^, ***η***_*m*_ = (*η*_*m*1_, *η*_*m*2_)^*T*^, and ***ξ***_*mg*_ = (*ξ*_*mg*1_, *ξ*_*mg*2_)^*T*^ . We also collect the pivot-adjusted growth rates 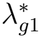 and 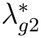 in a 2 × 2 diagonal matrix 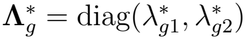. Re-writing equation (1) in vector form

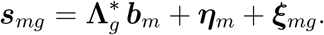

Eq. (1) describes fitness effect for a single mutation, however, to construct a JDFE, we need to specify fitness effects of all possible mutations. We generalize the approach of Reddy and Desai (2021) and Ardell et al. (2024), and assume that the slope of a given mutation is drawn from a bivariate distribution *p*_*sl*_(***b***_*m*_). For now, we do not assume a parametric form for *p*_*sl*_(***b***_*m*_).

Ardell et al. (2024) model the pivot noise *η* in a single environment (“pivot noise”) by a normal distribution. We generalize this by assuming a joint normal distribution *N* (***η***_*m*_; **0, Σ**_pi_) with mean **0** and a variance-covariance matrix given by:

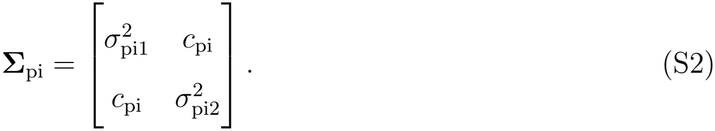

An intuitive understanding of this is as follows: since most global epistasis slopes are negative, mutations for negative adjusted growth rates in both environments are primarily beneficial and those for positive adjusted growth rates are primarily deleterious. Now, some mutations, those with a positive intercept in *η*_*me*_ *>* 0 will be beneficial specific to a given environment *e* even when adjusted growth rate in this environment is positive. If *c*_pi_ = 0, then these environment-specific beneficial mutations will be on an average neutral and deleterious for zero and positive adjusted growth rates. Similarly, if *c*_pi_ *>* 0, mutations that remain beneficial even after the typical mutation has become neutral or deleterious in environment *e* = 1 will statistically also remain beneficial in environment *e* = 2.

Idiosyncratic epistasis is also assumed to be joint normal with ***ξ***_*mg*_ *∼ N* (**0, Σ**_id_).

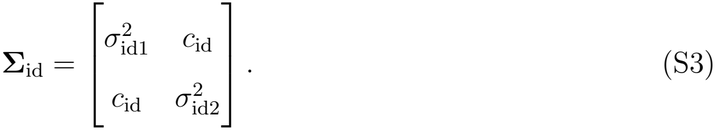

Note that this assumption differs from Reddy and Desai (2021), Ardell et al. (2024) and Brooks et al. (2026), in which the residuals depend on the slope of the mutation.

Further assuming that the effects of mutations are independent, conditional on its slope vector ***b***_*m*_, the fitness effect vector of mutation *m* in background strain *g* is distributed as 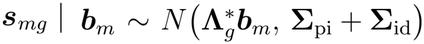. Therefore, the JDFE for strain *g* is obtained by marginalizing over all possible slope vectors:

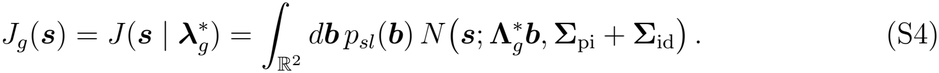

We can now use Eq. (S4) to derive statistics of the JDFE that can be used to test our theory against data and predict pleiotropic trajectories. As an illustrative example, consider the marginal mean fitness effect of a mutation in strain *g* and environment *e* = 1:

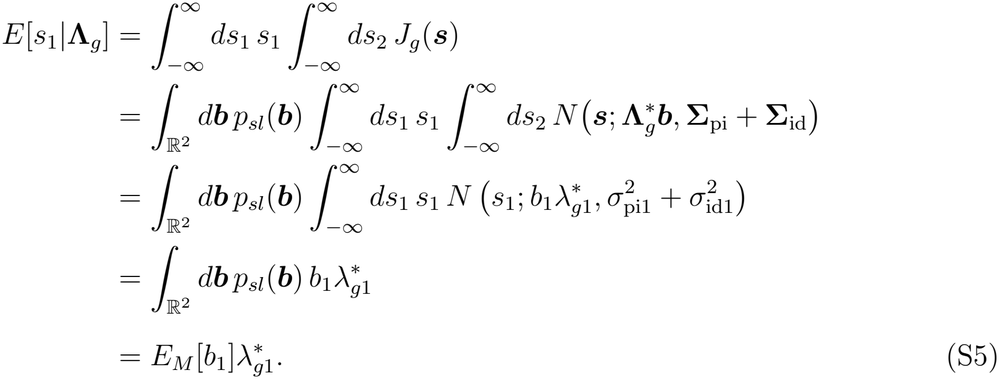

Similarly, 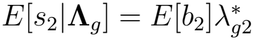, which implies that the expected fitness effect of a mutation changes linearly with the pivot-adjusted growth rate in each environment. These calculations recover those of Ardell et al. (2024), who derived the moments of the distribution of fitness effects (DFE).

#### S1.1.1 JDFE covariance

Next, we go beyond just marginal statistics and characterize properties of the joint fitness effects from the JDFE for a given strain *g*. We first calculate the covariance *Cov*(*s*_1_, *s*_2_) as:

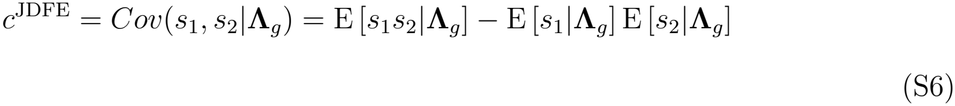

Since, E [*s*_1_|**Λ**_*g*_] and E [*s*_2_|**Λ**_*g*_] are known from Eq. (S5), we calculate E [*s*_1_*s*_2_|**Λ**_*g*_] first as:

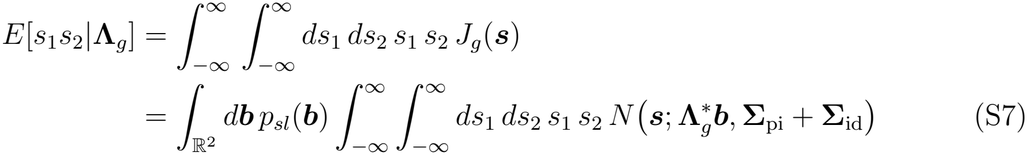

For a bivariate normal distribution, the cross-moment is equal to the product of the marginal means plus their covariance. Conditional on ***b***, the marginal means are 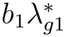 and 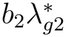, while covariance is *c*_pi_ + *c*_id_. Therefore the above expression simplifies to:

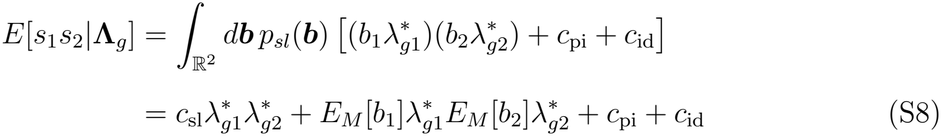

with *c*_sl_ = *Cov*(*b*_1_, *b*_2_) being the cross-environmental covariance between global epistasis slopes. Putting together Eq. (S6) and Eq (S8), we get the following:

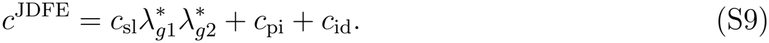

This equation tells us that the covariance *Cov*(*s*_1_, *s*_2_) of the JDFE changes linearly with the product of the pivot-adjusted growth rates 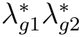 with a slope given by *c*_*sl*_ and intercept *c*_pi_ + *c*_id_.

#### S1.1.2 Pleiotropy statistic of the JDFE

When JDFEs do not depend on the background *g*, that is, there is no epistasis, the expected pleiotropic trajectory of fitness in environment *e* = 2 during adaptation to environment *e* = 1 only depends on the initial fitness 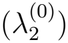 in *e* = 2 and changes linearly in time proportional to a background independent pleiotropy statistic *r*^JDFE^ (Ardell and Kryazhimskiy, 2021). The pleiotropy statistic is the expected fitness effect of a mutation in the non-home environment (here, *e* = 2) that is fixed in the home environment (here, *e* = 1).

However, when there is epistasis, the pleiotropy statistic will also depend on the background strain *g*, since fitness effects of mutations will now be background-dependent. More specifically, if there is global epistasis, pleiotropy statistic 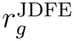 will depend on the background fitness in both environments (since the JDFE Eq. (S4) depends on background fitness). Further, for a given strain *g*, if 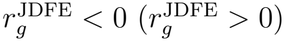, the expected fitness effect of a mutation will be negative (positive), causing a pleiotropic fitness loss (gain) in environment *e* = 2 during adaptation to environment *e* = 1. The magnitude of pleiotropic fitness gain and loss is also proportional to *r*^JDFE^. Thus, using this background fitness dependence, it is possible to construct pleiotropic trajectories.

Here, we derive 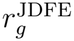 and a detailed analysis of pleiotropic trajectories is found in following sections:

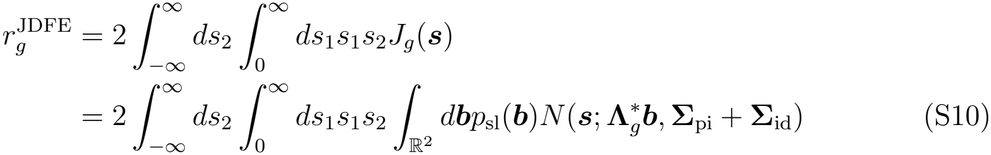

We can write the bivariate normal density 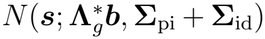 as the product of the conditional 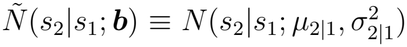 and marginal densities 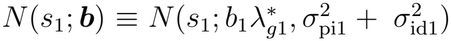 with,

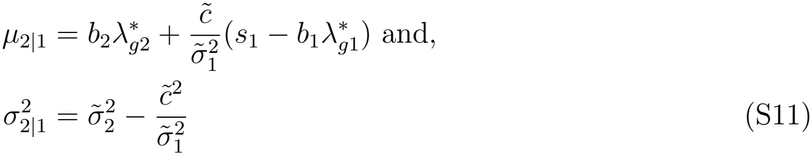

where 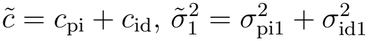 and 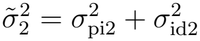. Thus, we get:

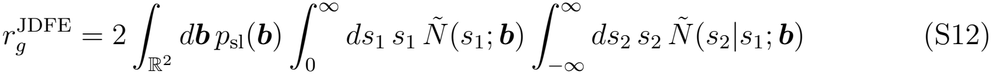

Plugging in the mean of the conditional distribution from Eq. (S11), this expression simplifies to,

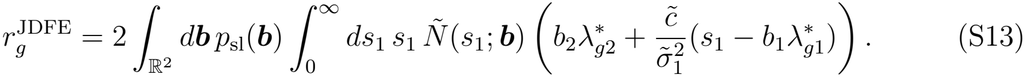

Expanding and rearranging, we get:

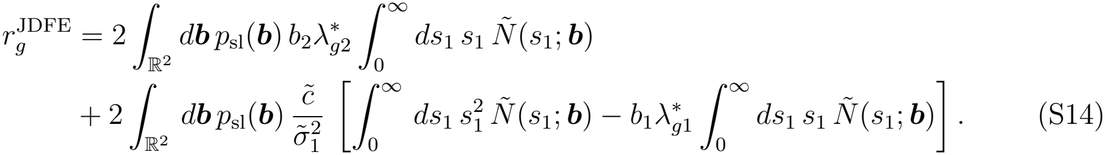

We can simplify this expression by using the moments of a normal distribution truncated to [0, ∞), and rearranging the general formula for a conditional expectation:

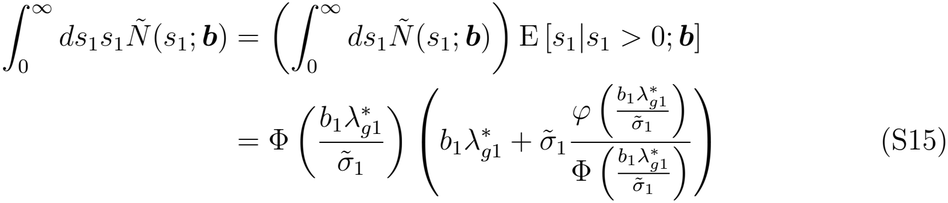

where *φ*(*z*) denotes the standard normal distribution and

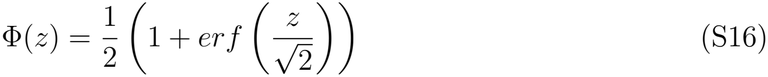

the standard normal CDF. Similarly, we have,

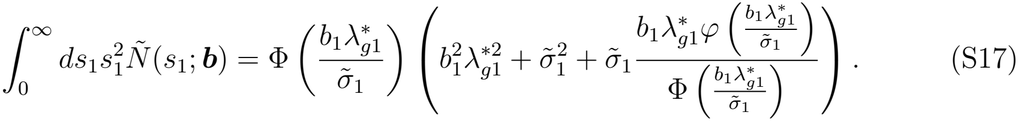

From equations (S15) and Eq. (S17), notice that,

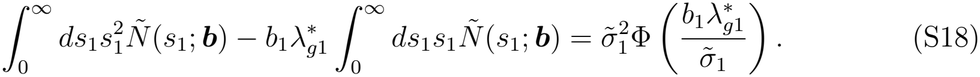

Using Eqs. (S15) and Eq. (S18), we now write *r*^JDFE^ in Eq. (S14) as:

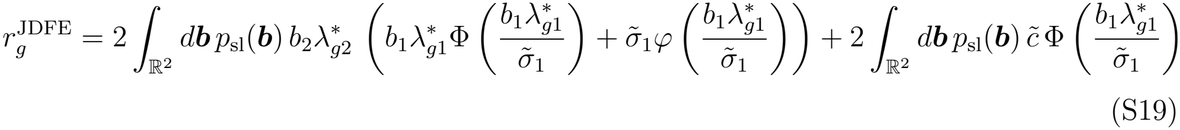

We write this equation in a simplified form as:

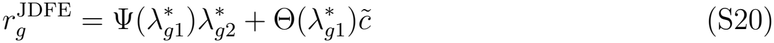

with 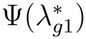 and 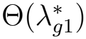 defined as in main text Eq. (10).

### S1.2 Predicted behavior of deterministic pleiotropic trajectories under SSWM

We will now discuss what the expression for pleiotropy statistic (Eq. (S20)) constrained by global epistasis implies for pleiotropic trajectories. First, however it is useful to also derive 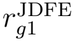, the quantity that governs adaptation to the first environment (Ardell et al., 2024) as:

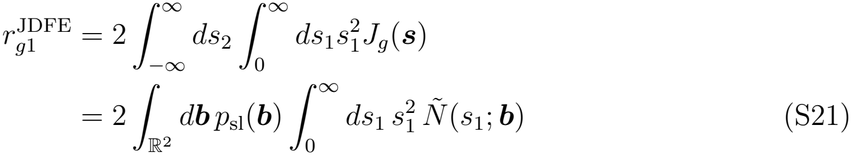

which from Eq. (S17) gives us:

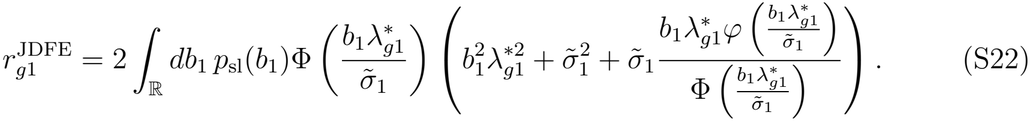

To derive the dynamics of the fitness trajectory, we use a deterministic approximation. Since *r*_1_ and *r*_2_ are the drift terms of the Fokker-Planck equation derived in Ardell and Kryazhim-skiy (2021), the expected mean fitness dynamics in both environments are given by the equations:

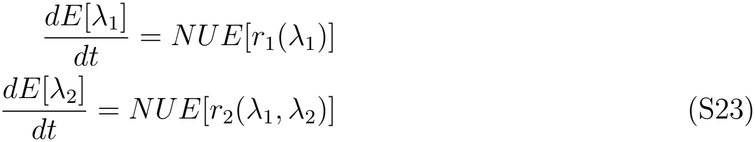

Here, *N* is the population size and *U* is the mutation rate. In general, *E*[*r*_*•*_(*λ*_*•*_)] ≠*r*_*•*_(*E*[*λ*_*•*_]), unless the function *r*_*•*_(*λ*_*•*_) is linear and the dynamics depend on higher order moments. However, we approximate the mean evolutionary trajectory by the deterministic path (see, for example, Aguadé-Gorgorió and Solé (2018)).

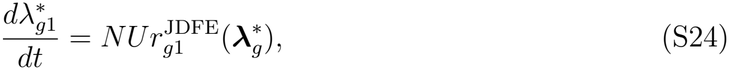

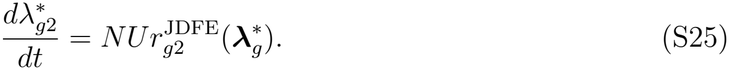

We do not formally check this approximation, rather we use the deterministic path to derive our basic predictions and check them against our simulations from a Gillespie algorithm. We derive predictions for pleiotropic trajectories in two distinct cases 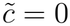 and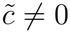.

#### S1.2.1 Case 1: Pivot noise and idiosyncratic terms are uncorrelated across environments

We first start with the case in which the pivot noise and idiosyncratic terms are both uncorrelated, that is 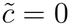. In this case,

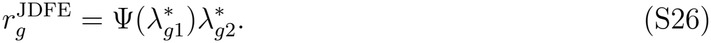

Decomposing the joint slope distribution *p*_sl_(***b***) into the conditional distribution *p*_sl_(*b*_2_|*b*_1_) and the marginal *p*_sl_(*b*_1_), we can simplify the above parameters as:

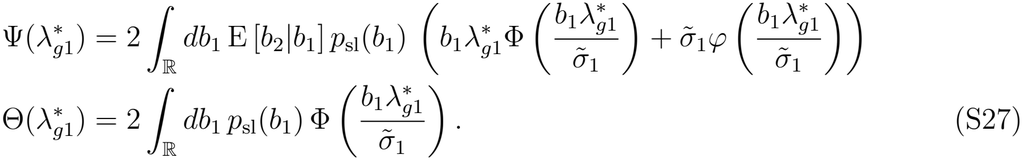

Using the truncated moment expression in Eq. (S15) we first rewrite the expression in Eq. (S27) as:

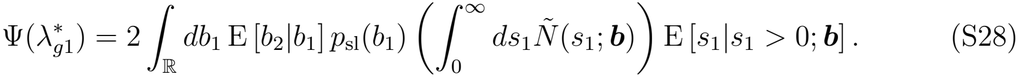

Since 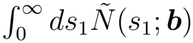 is a cumulative density, it is positive and E [*s*_1_|*s*_1_ *>* 0; ***b***] is an expectation over only positive values, so it is also positive. Thus, as long as the expectation of the global epistasis slope in environment 2 conditioned on the slope in environment 1, E [*b*_2_|*b*_1_], is negative for all slopes, 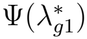 is negative. We assume below that this is the case, since empirically, most slopes are negative and indistinguishable across environments.

With the assumption that E [*b*_2_|*b*_1_] *<* 0, for all values of 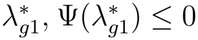. As a consequence, we have 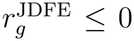 whenever 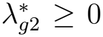, and 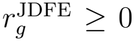, whenever 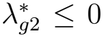 . Thus, if fitness in environment 2 lies below the pivot 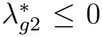, pleiotropic fitness gains are expected and if it is above 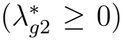, pleiotropic fitness losses are expected. Additionally, since 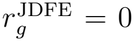 whenever 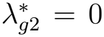, the pivot growth rate corresponding to 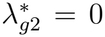 is an attractor for the deterministic dynamics.

To sketch out the dynamics of pleiotropic changes, we use some asymptotics in addition to the above analysis. During adaptation, by definition, fitness in environment 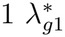 increases. Thus, we consider three regimes corresponding to early 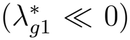, mid 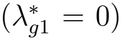 and late adaptation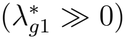, assuming fitness starts way below the pivot in environment 1. When 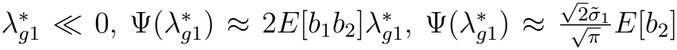 when 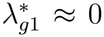 and 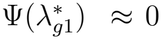 when 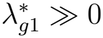. Therefore, we expect that 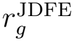 will decrease with a steeper slope when 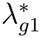 is highly negative and saturate at zero as 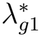 increases. Thus, if the initial fitness in environment 2 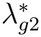 is below (above) the pivot in environment 2, when fitness is low in environment 1, there will be rapid pleiotropic fitness gains (losses) that slow and potentially converge to the pivot attractor. We test this prediction in simulations.

#### S1.2.2 Case 2: Pivot noise and idiosyncratic terms are correlated across environments

Unlike case 1, the pivot fitness in environment 2 is not an attractor, instead, when 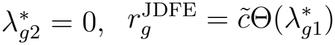, which generally depends on 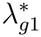. We will now show that in the special case that the slopes *b*_1_ = *b*_2_ = *b*, similar to what is observed in our dataset, there is an attracting line toward which trajectories converge given by main text Eq. (5). We will first note that the ratio of instantaneous rates of change of 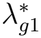 and 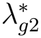 are given by the ratio of the pleiotropy and adaptation statistic.

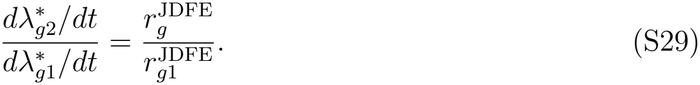

To obtain a relationship between the two statistics, we first rearrange Eq. (S14) as follows:

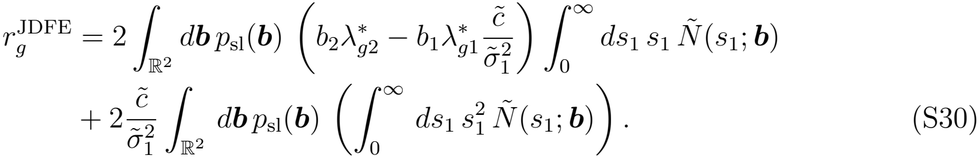

By definition of the adaptation statistic Eq. (S21), the above expression is written as:

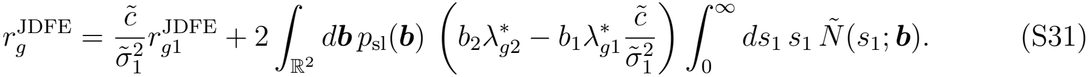

Now suppose that the slopes remain invariant across environments. In other words, *b*_1_ = *b*_2_ = *b* and *p*_sl_(*b*) is the slope distribution, we get:

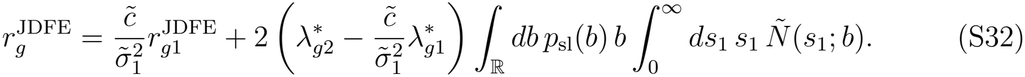

In the special case that Eq. (5) is true, the above expression simplifies to:

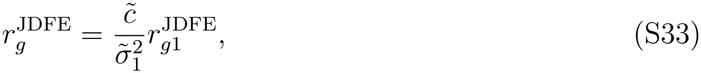

which can also be written as:

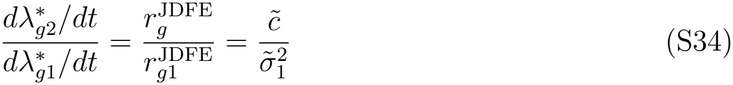

which integrates to:

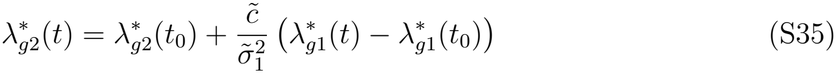

This satisfies Eq. (5) when 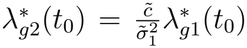. Thus, a trajectory that starts on the line defined by Eq. (5) remains on it.

To now show that the line defined by Eq. (5) is attracting in the case of same slopes, consider a perturbation on the trajectory:

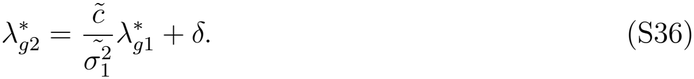

Differentiating with respect to time, we get,

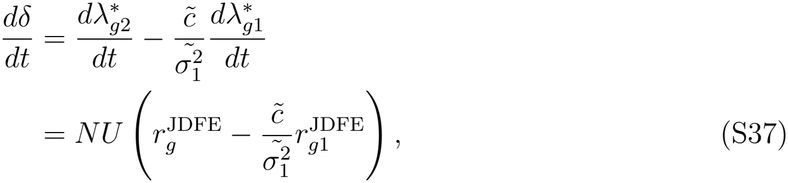

where the constant *NU >* 0. From Eq. (S32), we get,

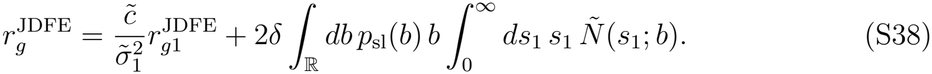

Combining Eq. (S38) and Eq. (S37)

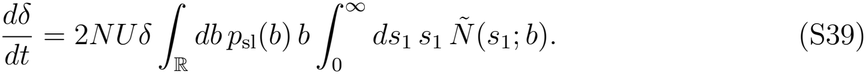

Now, 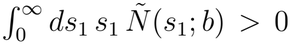 regardless of the value of 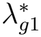, which means that if all slopes *b <* 0,

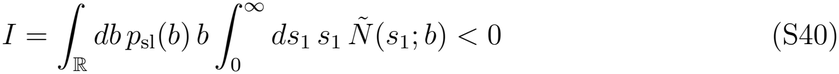

Therefore, if 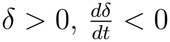 and if 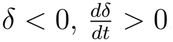. Thus, perturbations from the line defined in Eq. (5) are drawn closer to it, which implies it is attracting.

### S1.3 Implementation of LMM

We describe here the formulas used to implement our modified linear mixed effects model in `glmmTMB`. The estimate for the fitness effect of a mutation labeled `Mut_ID` (*m*) in strain labeled `Strain` (*g*) in environment env (*e*) for biological replicate `bioRep` (*r*) is denoted by `avgS` (*s*_*mgre*_). The unadjusted and adjusted growth rates (fitness) are denoted by `Mean_GR` (*λ*_*gre*_) and `adjusted_GR` 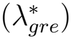, respectively. Additionally, we define `one_plus_slope` as a quantity that stores the predicted slope per mutation plus 1 (and environment for different slopes model).

We initially compare the following two models to test whether the slopes and intercepts are negatively correlated for the unadjusted fitness like found in Ardell et al. (2024). To do so, we go through all environments separately and fit the following models.

~~~
MP: avgS ∼ 1 + Mean_GR + (1 + Mean_GR | Mut_ID)
~~~

and,

~~~
MP0: avgS ∼ 1 + Mean_GR + (1 | Mut_ID) + (0 + Mean_GR | Mut_ID).
~~~

We compare these two models using a likelihood ratio test. `MP` accounts for slope intercept correlation and `MP0` does not. Since the correlations are significant (see Supplementary data), we estimate a common environment-specific pivot for both the same slope and different slope models.

We estimate a common pivot fitness using the regression coefficient obtained from the following model which is fit across environments unlike `MP`, which is fit per environment.

~~~
MP1: avgS ∼ 1 + Mean_GR + (0 + Mean_GR + env | Mut_ID)
~~~

We use this pivot estimate for both the same slope and different slope models. Using the pivot, we calculate adjusted fitness of all strains for each replicate and environment and fit the following preliminary model to get global epistasis slopes, denoting the same slope model as `MSS0` and different slopes model as `MDS0`, we have

~~~
MSS0: avgS ∼ 0 + adjusted_GR + (0 + adjusted_GR| Mut_ID)
+ (0 + env | Mut_ID)
MDS0: avgS ∼ 0 + adjusted_GR + (0 + adjusted_GR:env| Mut_ID)
+ (0 + env | Mut_ID)
~~~

Using these preliminary model-predicted slopes we fit the full model in two iterations to ensure that we obtain similar estimates each time:

~~~
MSS: avgS ∼ 0 + adjusted_GR + (0 + adjusted_GR| Mut_ID)
+ ( 0 + env | Mut_ID)+ (0 + env | Mut_ID:Strain) +
diag(0 + one_plus_slope:env|bioRep:Strain)+ diag(0 + env|Mut_ID:bioRep:Strain)
~~~

~~~
MDS: avgS ∼0 + adjusted_GR:env + (0 + adjusted_GR:env| Mut_ID)
+ ( 0 + env | Mut_ID)+ (0 + env | Mut_ID:Strain)
+ diag(0 + one_plus_slope:env|bioRep:Strain)+ diag(0 + env|Mut_ID:bioRep:Strain)
~~~

## S2 SI Figures

**Figure S1.**
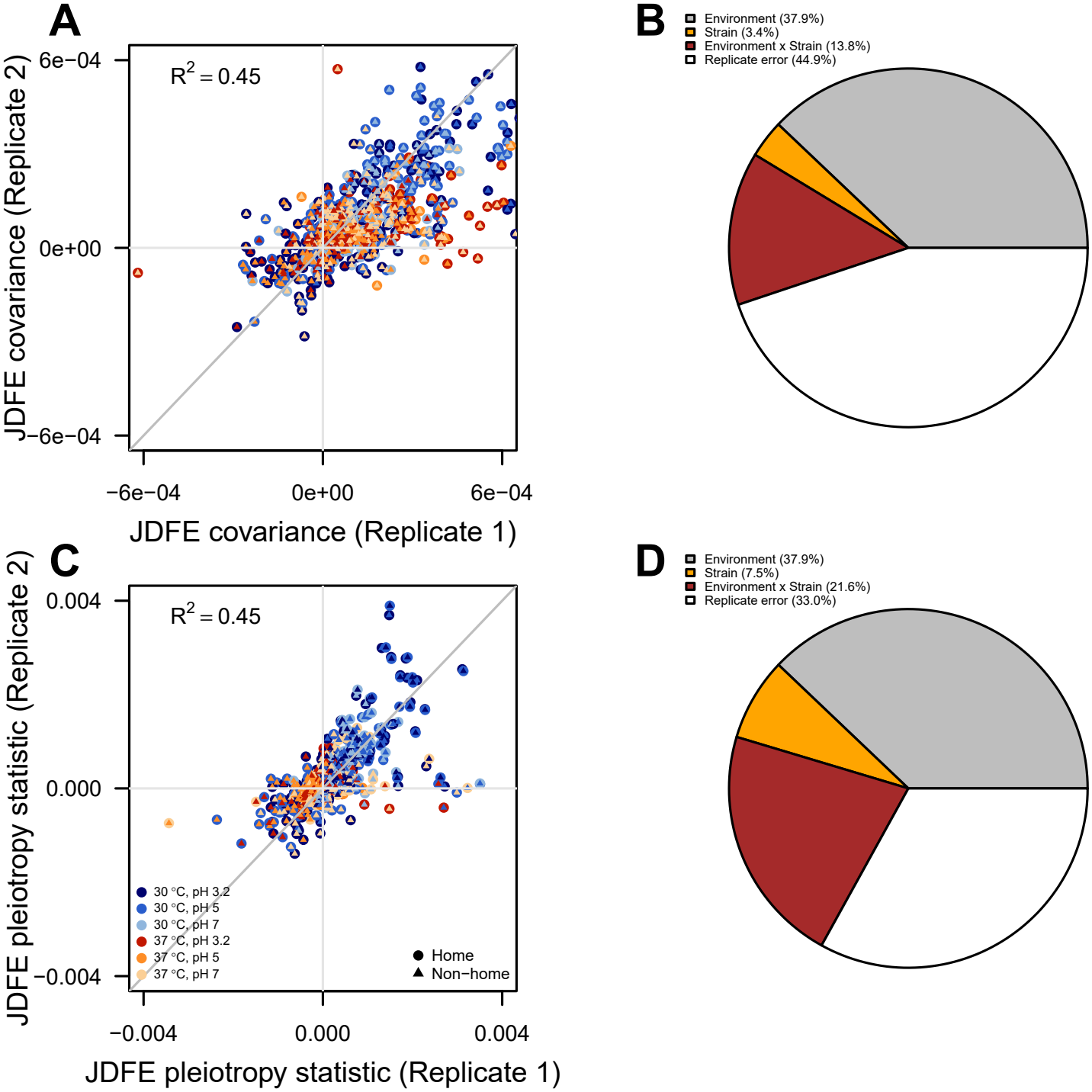
Estimates of JDFE covariance (A) and the pleiotropy statistic (C) from two biological replicates plotted against each other for all strains and environment pairs. Each point represents a strain-environment-pair combination. Filled circles indicate the color of the home environment, while overlaid triangles indicate the color of the non-home environment. Partition of variation in JDFE covariance (B) and the pleiotropy statistic (D) into components associated with environment pair, strain, environment pair-strain interaction, and residual replicate-level variation. Variance components were estimated using linear mixed-effects models with environment pair as a fixed effect and strain and environment-pair-by-strain interaction as random effects.

**Figure S2.**
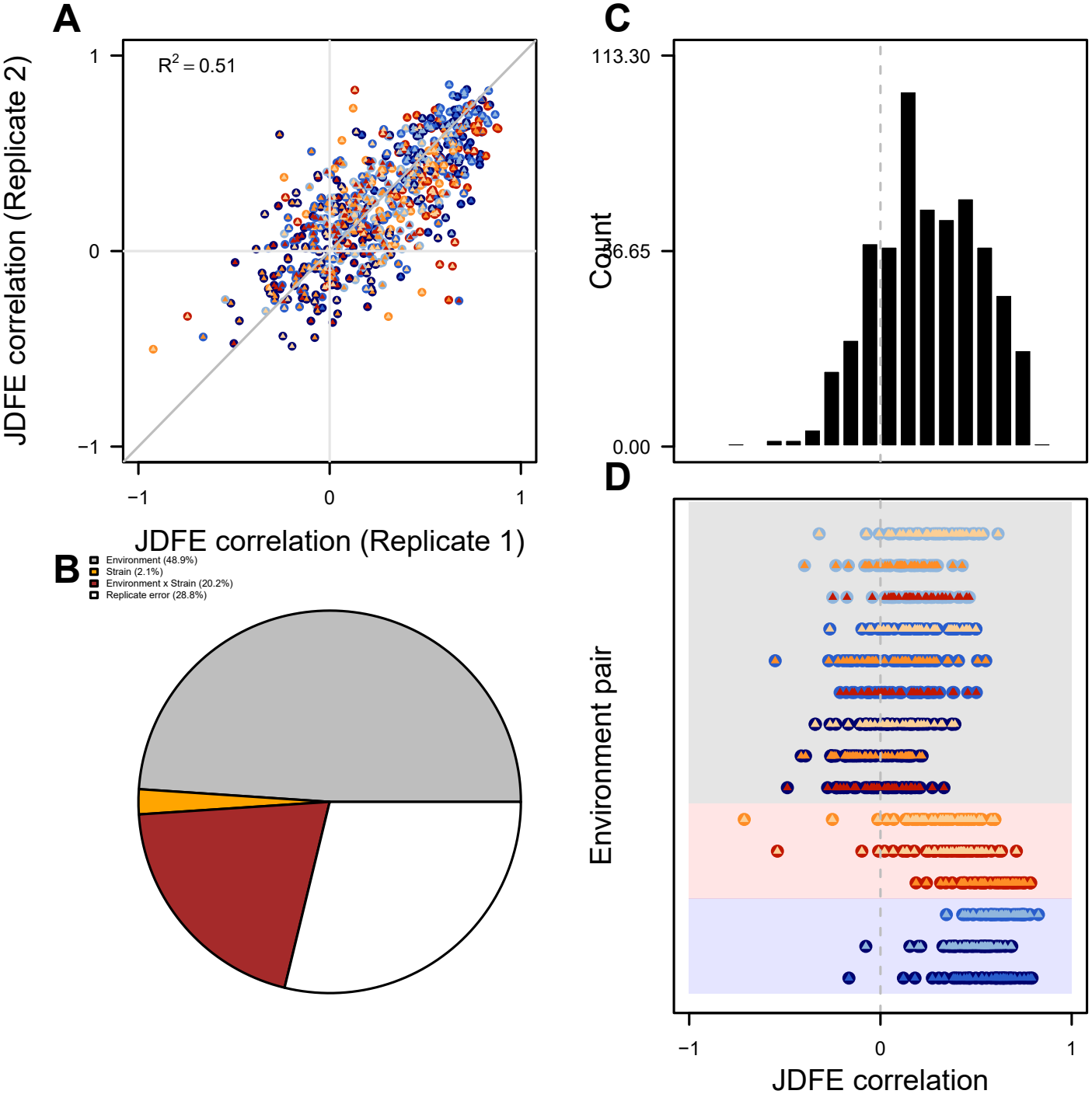
A: Estimates of JDFE correlation for two biological replicates plotted against each other for all strains and environment pairs. Each point represents a strain-environment pair combination. B: Partition of variation in JDFE correlation into components associated with environment pair, strain, environment pair-strain interaction, and residual replicate-level variation. Variance components were estimated using a linear mixed-effects model with environment pair as a fixed effect and strain and environment pair-strain interaction as random effects. C: Distribution of JDFE correlation across strains and environment pairs, averaged over the two biological replicates. D: Replicate-averaged JDFE correlation grouped by environment pair. Each point represents one strain-environment pair combination.

**Figure S3.**
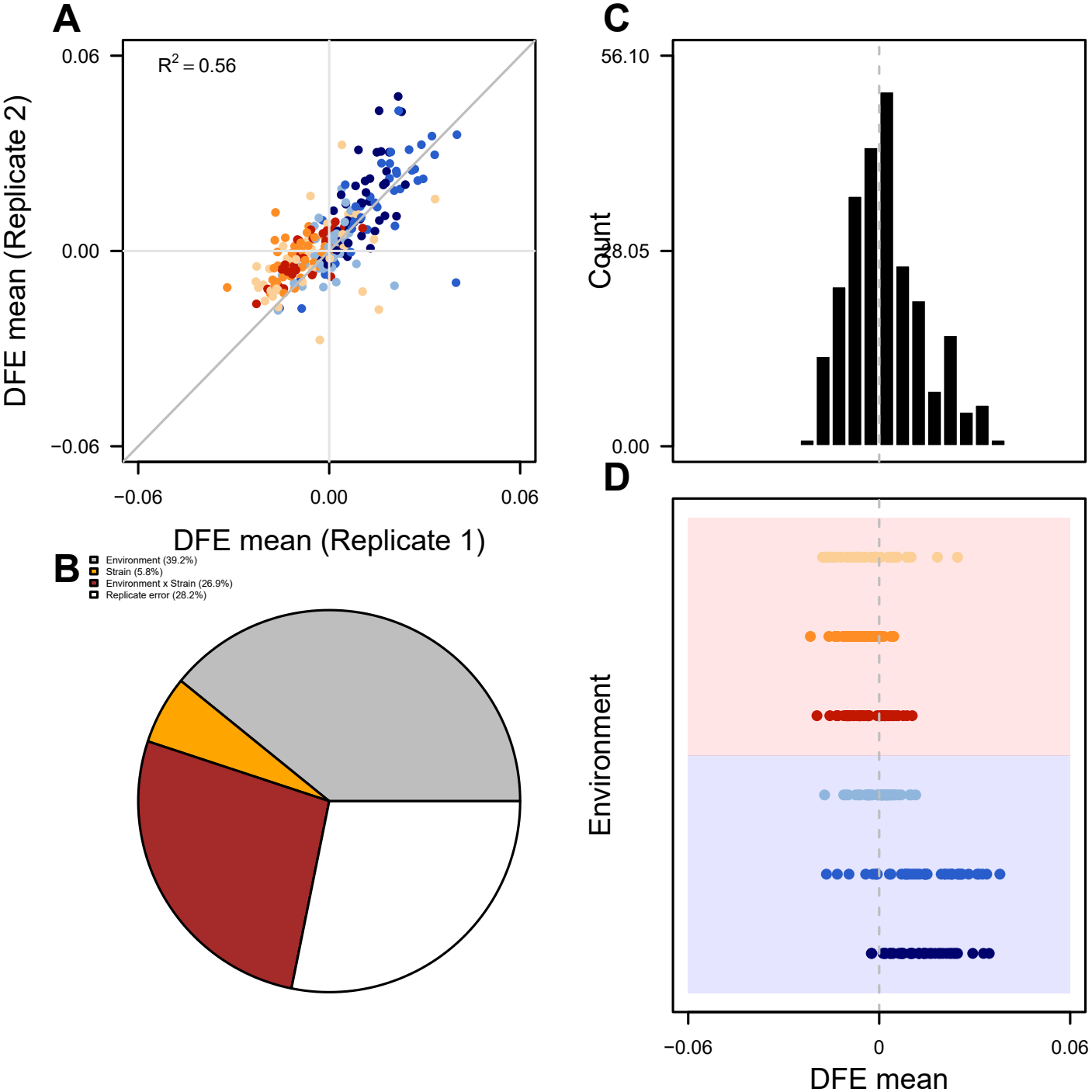
A: Estimates of marginal DFE mean for two biological replicates plotted against each other for all strains and environments. Each point represents a strain-environment combination. B: Partition of variation in DFE mean into components associated with environment, strain, environment-by-strain interaction, and residual replicate-level variation. Variance components were estimated using a linear mixed-effects model with environment as a fixed effect and strain and environment-strain interaction as random effects. C: Distribution of DFE means across strains and environments, averaged over the two biological replicates. D: Replicate-averaged DFE means grouped by environment. Each point represents one strain-environment combination.

**Figure S4.**
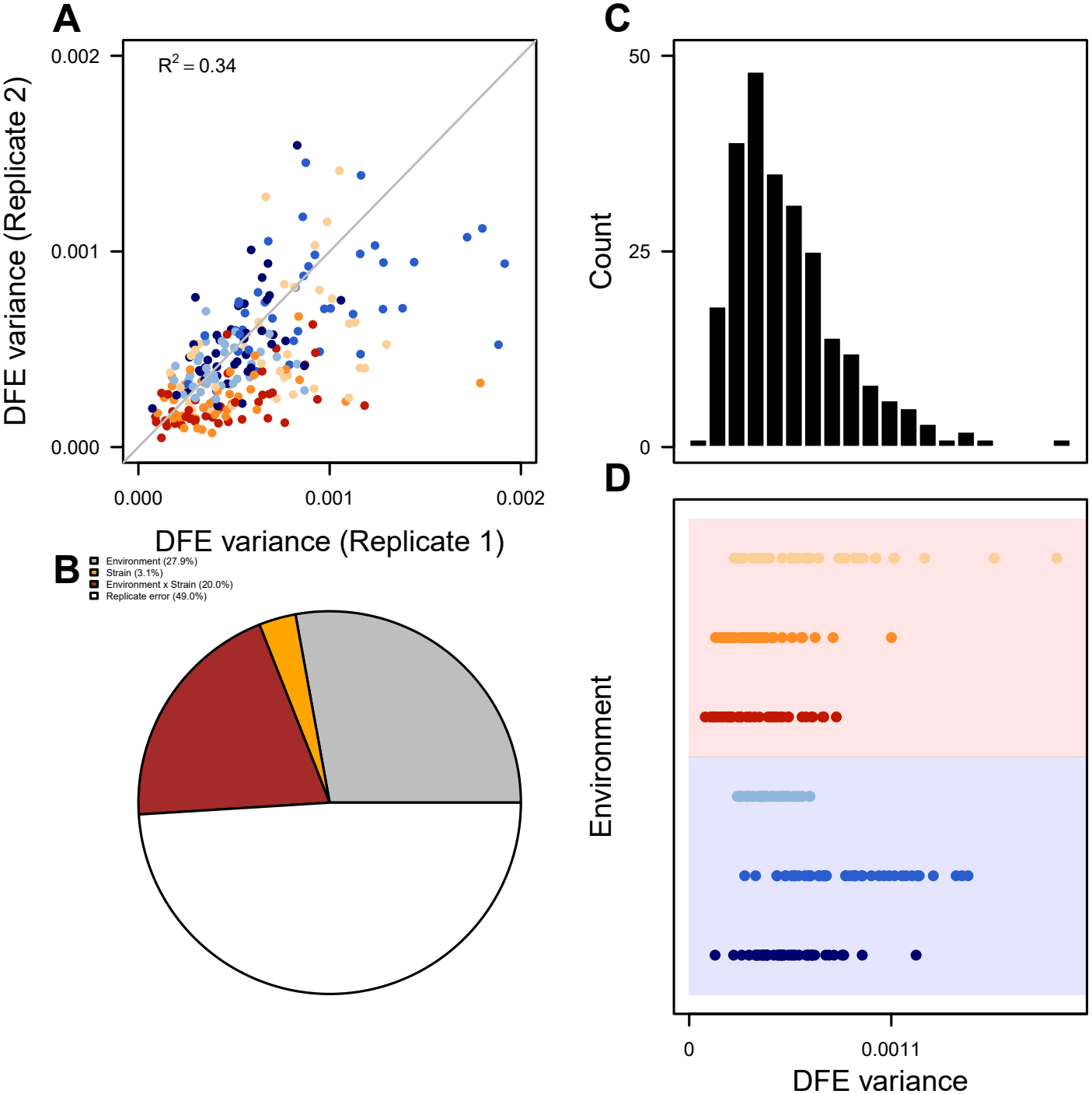
A: Estimates of marginal DFE variance for two biological replicates plotted against each other for all strains and environments. Each point represents a strain-environment combination. B: Partition of variation in DFE variance into components associated with environment, strain, environment-by-strain interaction, and residual replicate-level variation. Variance components were estimated using a linear mixed-effects model with environment as a fixed effect and strain and environment-strain interaction as random effects. C: Distribution of DFE variances across strains and environments, averaged over the two biological replicates. D: Replicate-averaged DFE variances grouped by environment. Each point represents one strain-environment combination.

**Figure S5.**
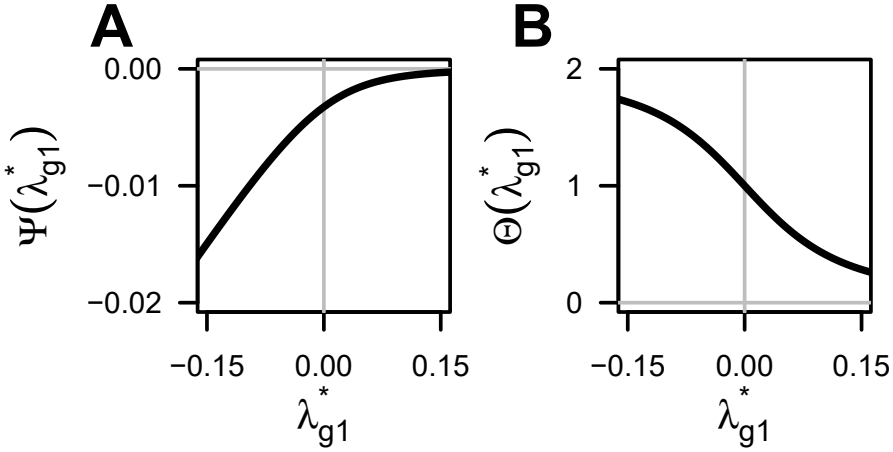
A: Function 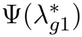 and B: 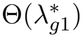. Functions were evaluated numerically using the same slopes model, with the mean global epistasis slope parameters and mean remaining standard deviation across the six environments. The plotted range of adjusted home fitness corresponds to that observed in the Ardell24 dataset.

**Figure S6.**
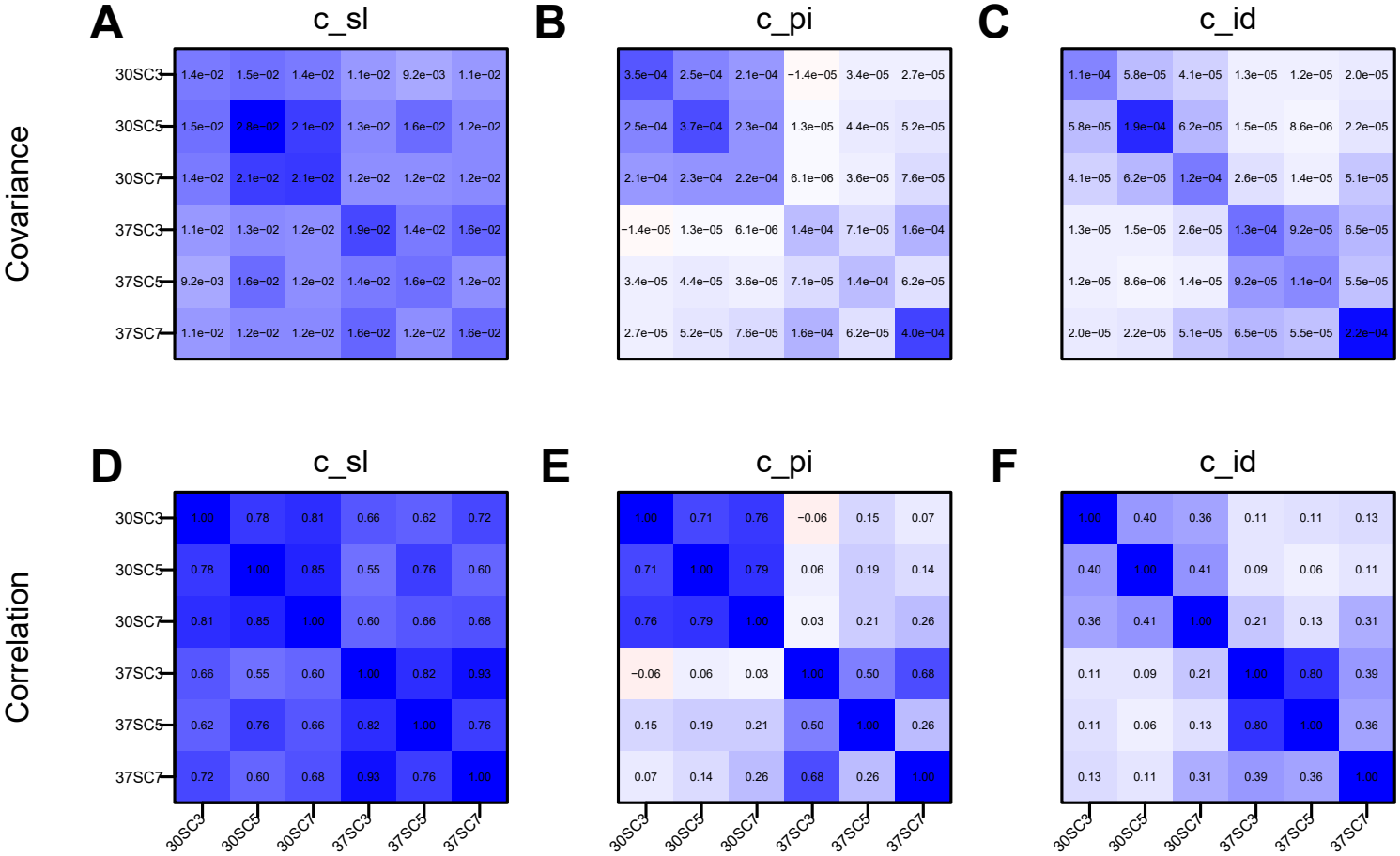
Estimated covariance and correlation matrices from the different slopes model in which GE slopes can vary across environments for a given mutation. A–C: Cross-environment covariance matrices for global epistasis slopes (*c*_sl_), pivot noise (*c*_pi_), and idiosyncratic epistasis (*c*_id_), respectively. D–F: Corresponding correlation matrices. Rows and columns represent the six environments, varying in temperature (30^*°*^C and 37^*°*^C) and pH (3.2, 5.0, and 7.0). Pivot noise and idiosyncratic correlations are generally stronger between environments at the same temperature than between environments at different temperatures.

**Figure S7.**
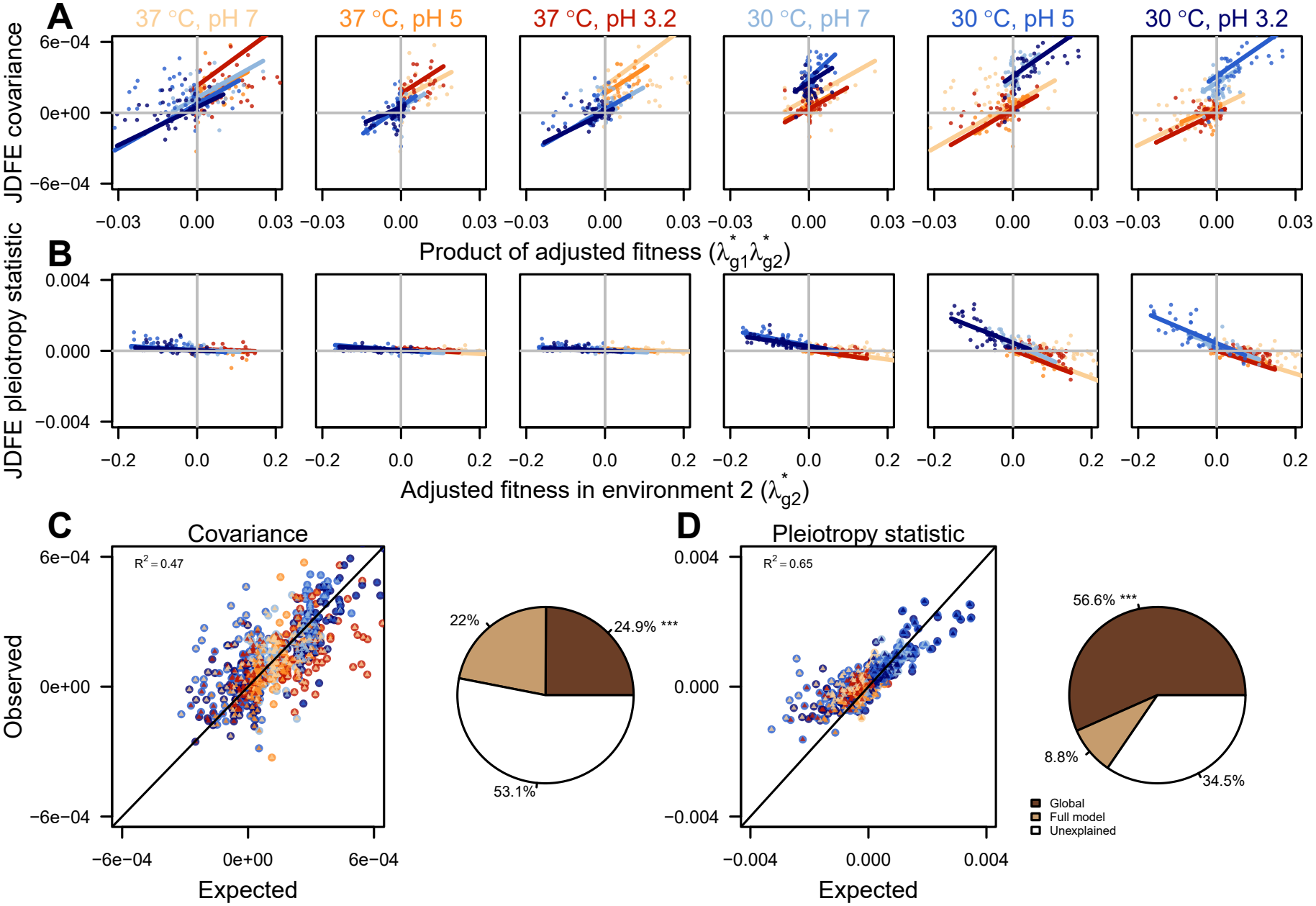
Different slopes model and same slope model (Main text Figure 2) are comparable. A: JDFE covariance vs. product of adjusted fitness in environments 1 and 2. Each panel corresponds to one home environment. Points represent individual strain-environment-pair JDFEs, and solid lines show theoretical predictions from the different slopes global epistasis model. B: Pleiotropy statistic vs. fitness in environment 2. Solid lines show theoretical predictions evaluated at the median adjusted background fitness in environment 1 for the corresponding environment pair. C: Observed vs. expected JDFE covariance and variation partitioned into global epistasis and the remaining model terms. D: Observed vs. expected JDFE pleiotropy statistic and variation partitioned into global epistasis and the remaining model terms.

**Figure S8.**
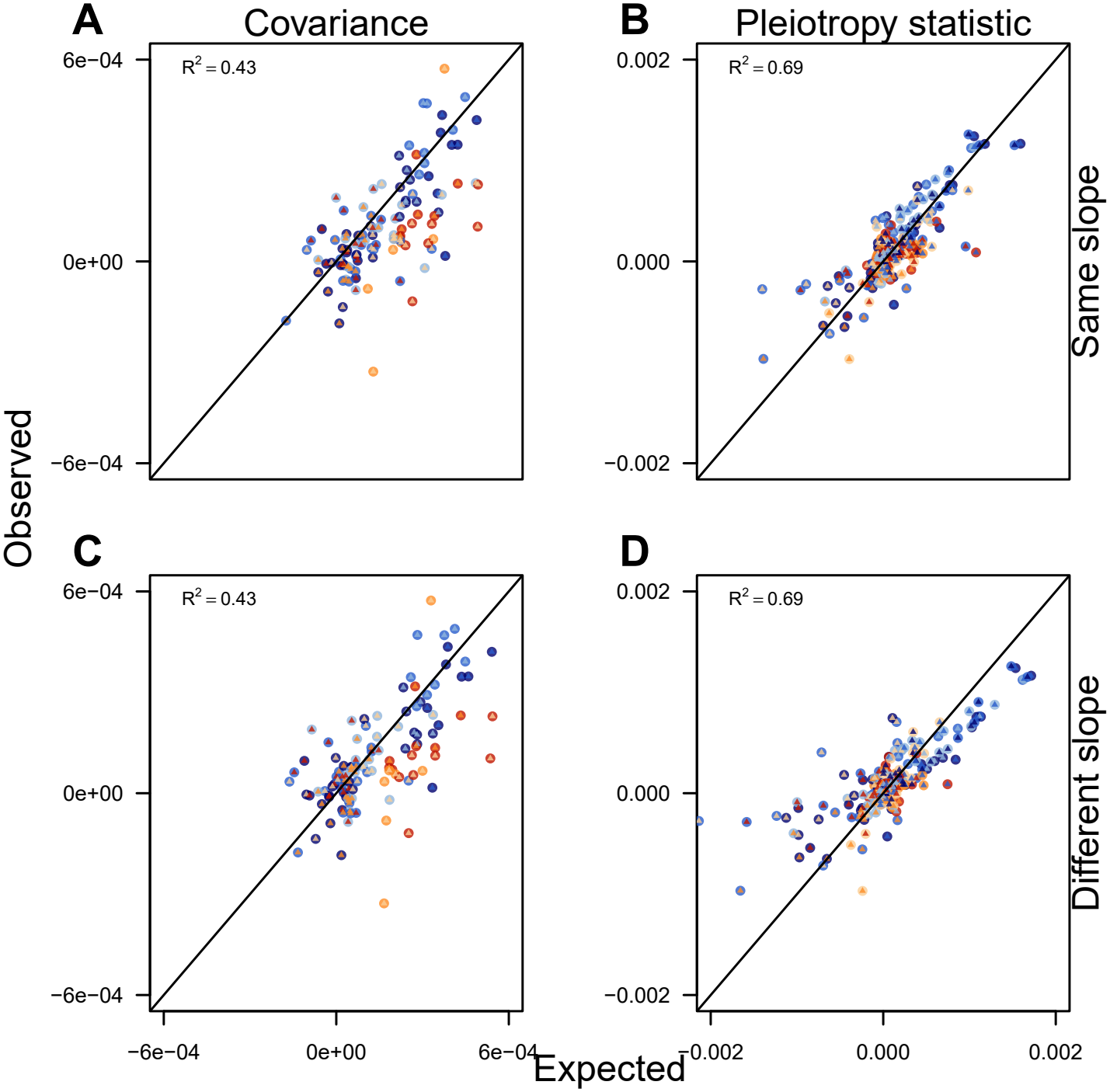
Same slope and different slope LMMs in Methods section 4.2.2 are fit using only 34 “training” strains (80 percent) and predictions are made for 8 test strains across environment pairs. For both models (A–B same slope and C–D different slope), the LMM predicted and observed JDFE covariance (A and C) and pleiotropy statistic (B and D) agree with each other.

**Figure S9.**
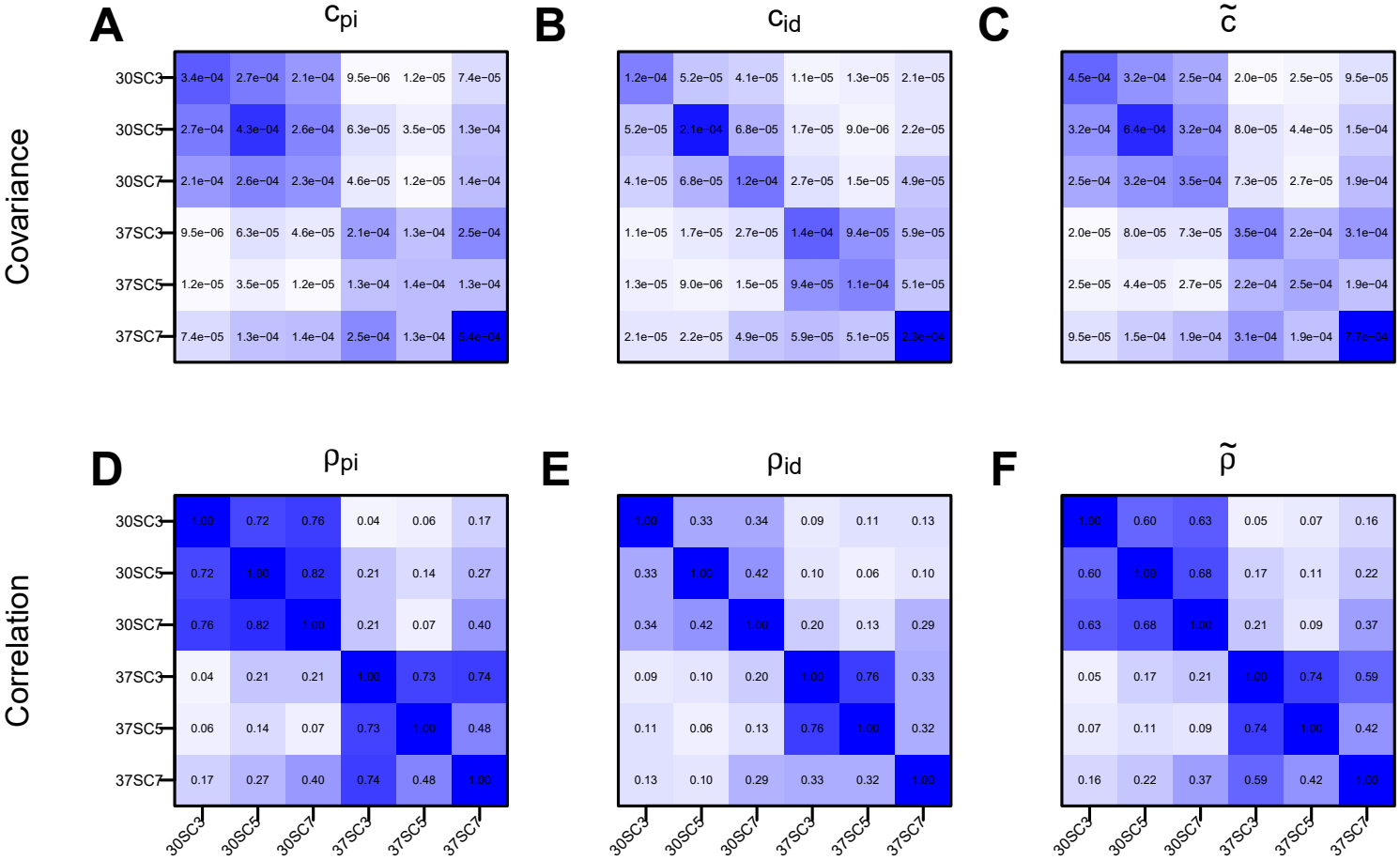
Estimated covariance and correlation matrices from the same slope model. A–C Cross-environment covariance matrices for pivot noise (*c*_pi_), idiosyncratic epistasis (*c*_id_) and pleiotropic covariance 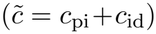, respectively. D–F: Corresponding correlation matrices. Rows and columns represent the six environments, varying in temperature (30^*°*^C and 37^*°*^C) and pH (3.2, 5.0, and 7.0). Pivot noise and idiosyncratic correlations are generally stronger between environments at the same temperature than between environments at different temperatures.

**Figure S10.**
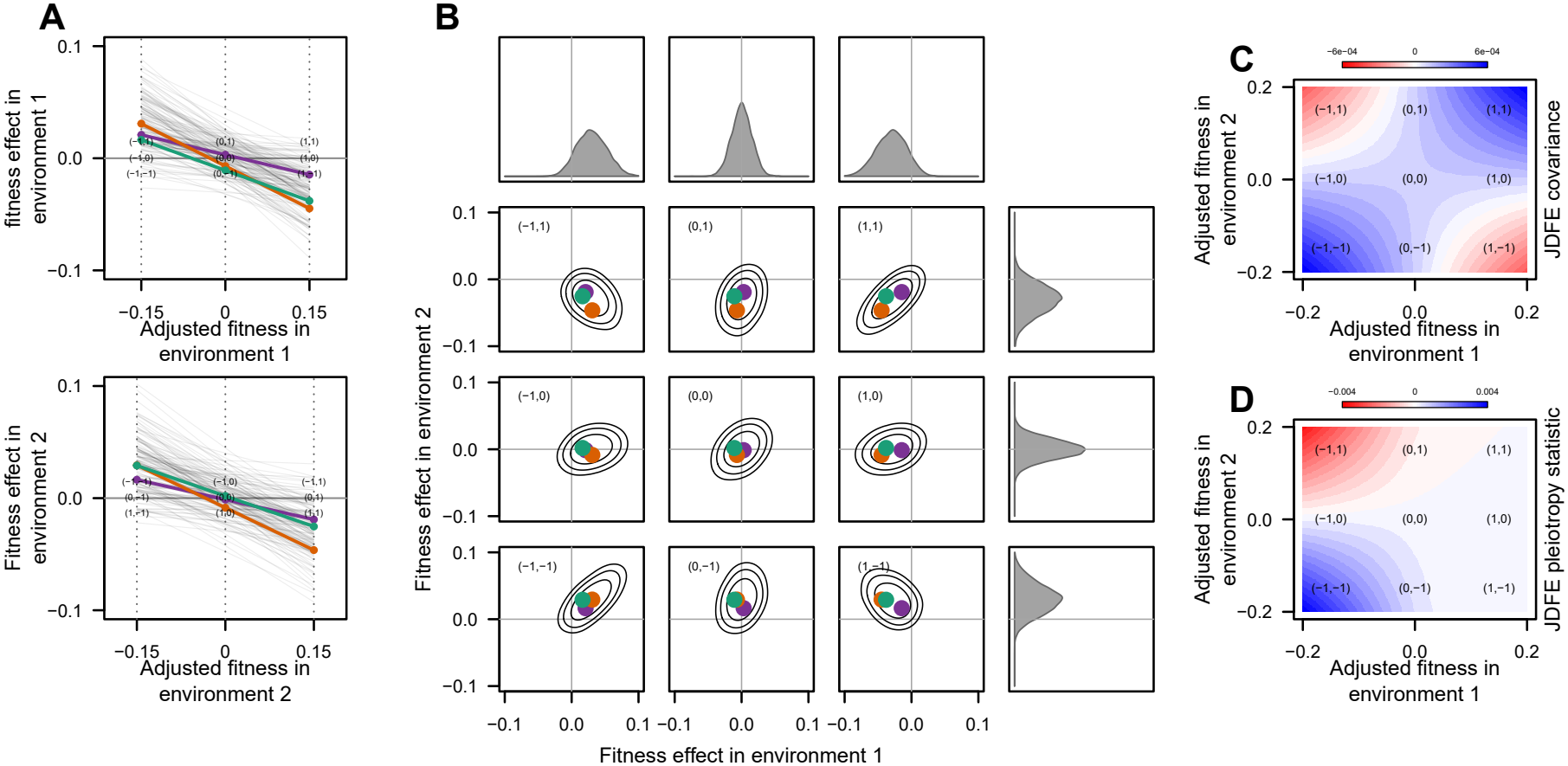
Predicted JDFE and properties for same slope and positive pleiotropic covariance. A : Illustrative fitness effects vs. fitness plots for mutations exhibiting global epistasis with the same slope with positive pivot covariance in environments 1 (top row) and 2 (bottom row), with three mutations highlighted. We additionally highlight nine strains with different combinations of adjusted fitness relative to pivot fitness. Labels (−1, 0, 1) denote adjusted fitness below, at, or above the pivot, respectively. For example, (-1,-1) indicates that strain fitness is below the pivot in environments 1 and 2. B: JDFE and marginal DFEs for the highlighted strains. C: Predicted JDFE covariance and D: pleiotropy statistic for different combinations of adjusted fitness in environments 1 and 2.

**Figure S11.**
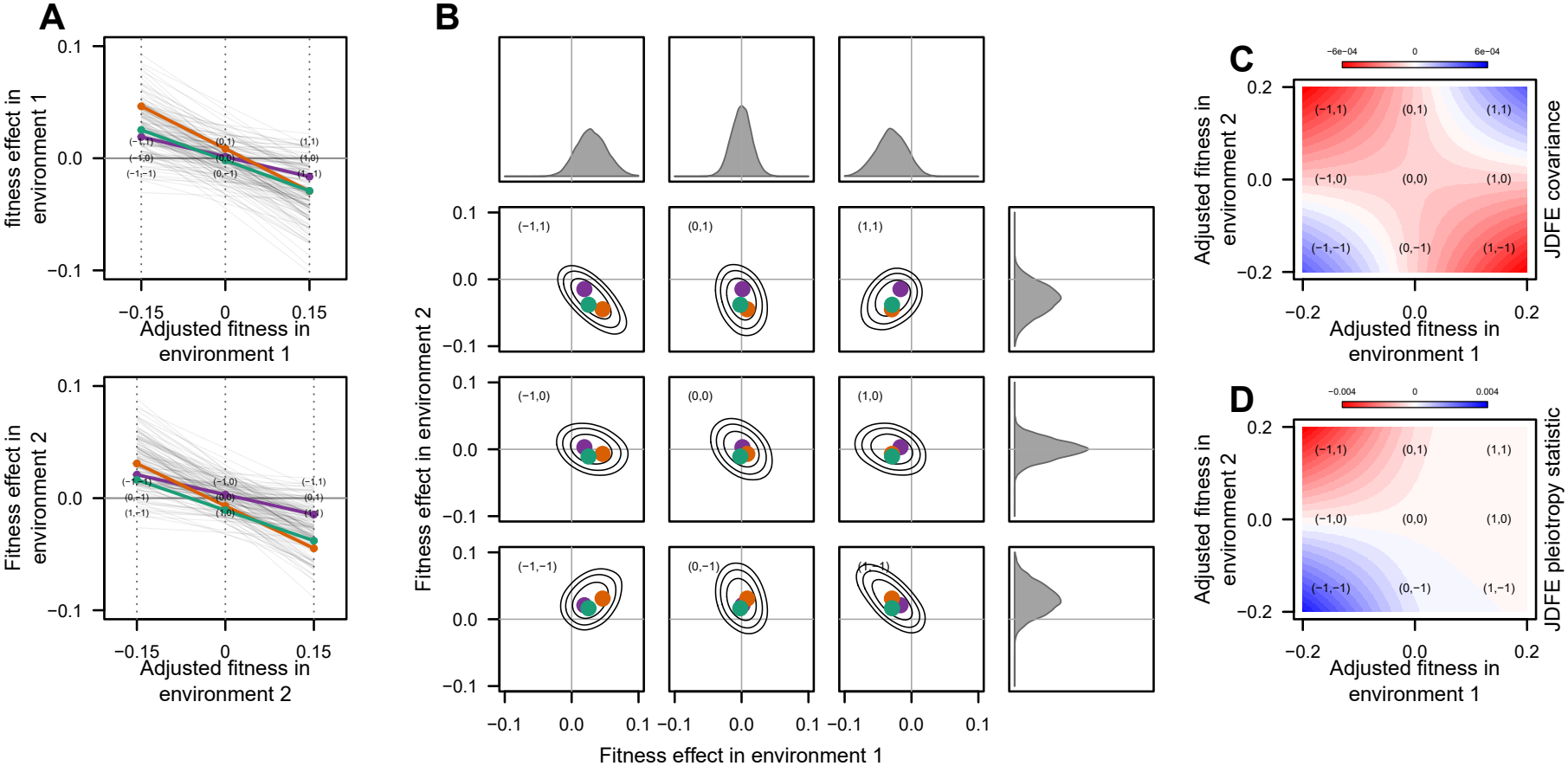
Predicted JDFE and properties for same slope and negative pleiotropic covariance. A : Illustrative fitness effects vs. fitness plots for mutations exhibiting global epistasis with the same slope with negative pivot covariance in environments 1 (top row) and 2 (bottom row), with three mutations highlighted. We additionally highlight nine strains with different combinations of adjusted fitness relative to pivot fitness. Labels (−1, 0, 1) denote adjusted fitness below, at, or above the pivot, respectively. For example, (-1,-1) indicates that strain fitness is below the pivot in environments 1 and 2. B: JDFE and marginal DFEs for the highlighted strains. C: Predicted JDFE covariance and D: pleiotropy statistic for different combinations of adjusted fitness in environments 1 and 2.

**Figure S12.**
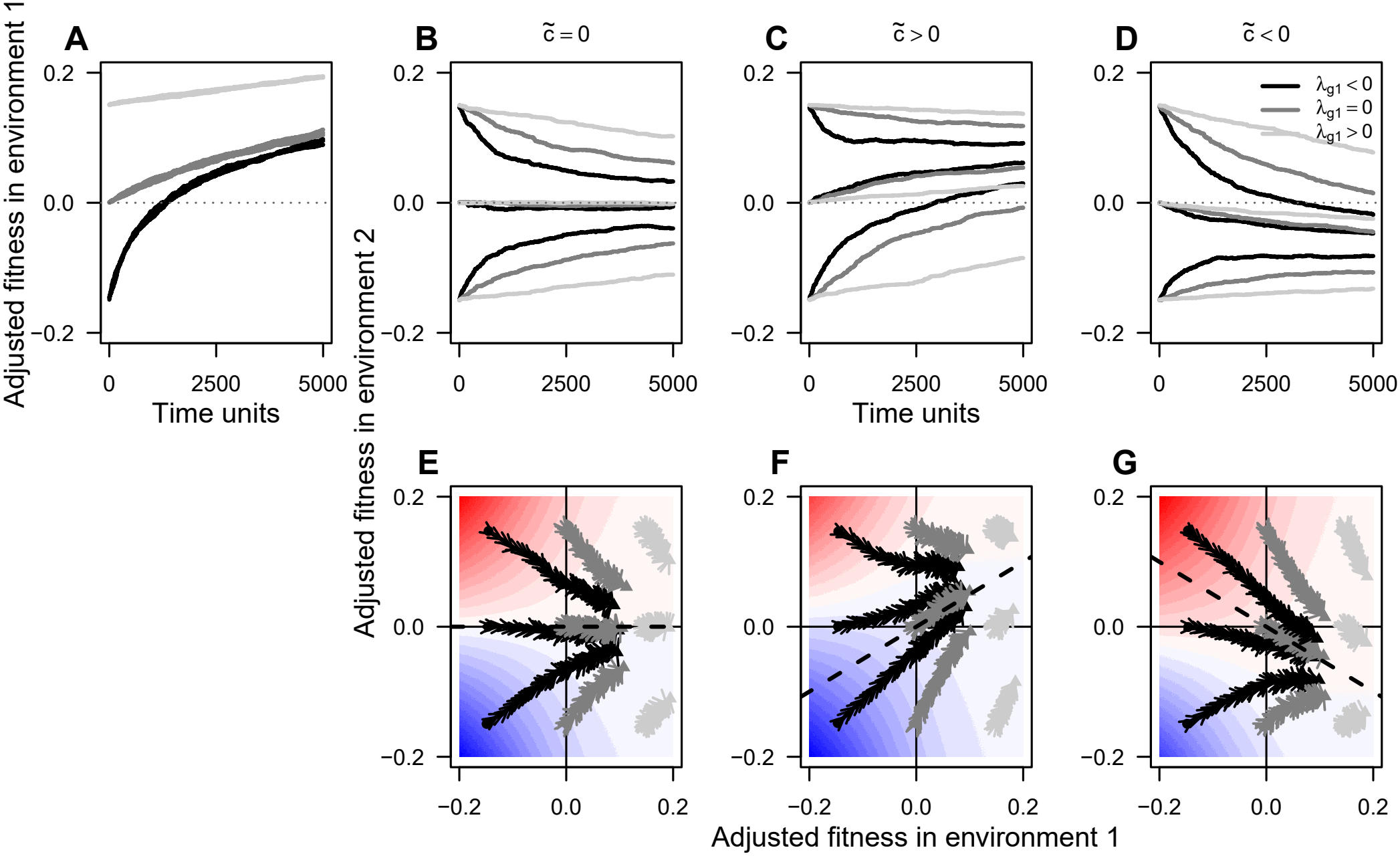
Predicted trajectories when global epistasis slopes are uncorrelated. Same parameters are used as main text Figure 4, except there is no correlation between global epistasis slopes *ρ*_sl_ = 0. For no pleiotropic covariance 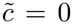, as predicted by our theory, the trajectories converge to the pivot fitness. For 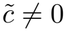 cases, the dashed line is no longer the attractor, nevertheless, the qualitative dynamics are similar to the same slope case.

**Figure S13.**
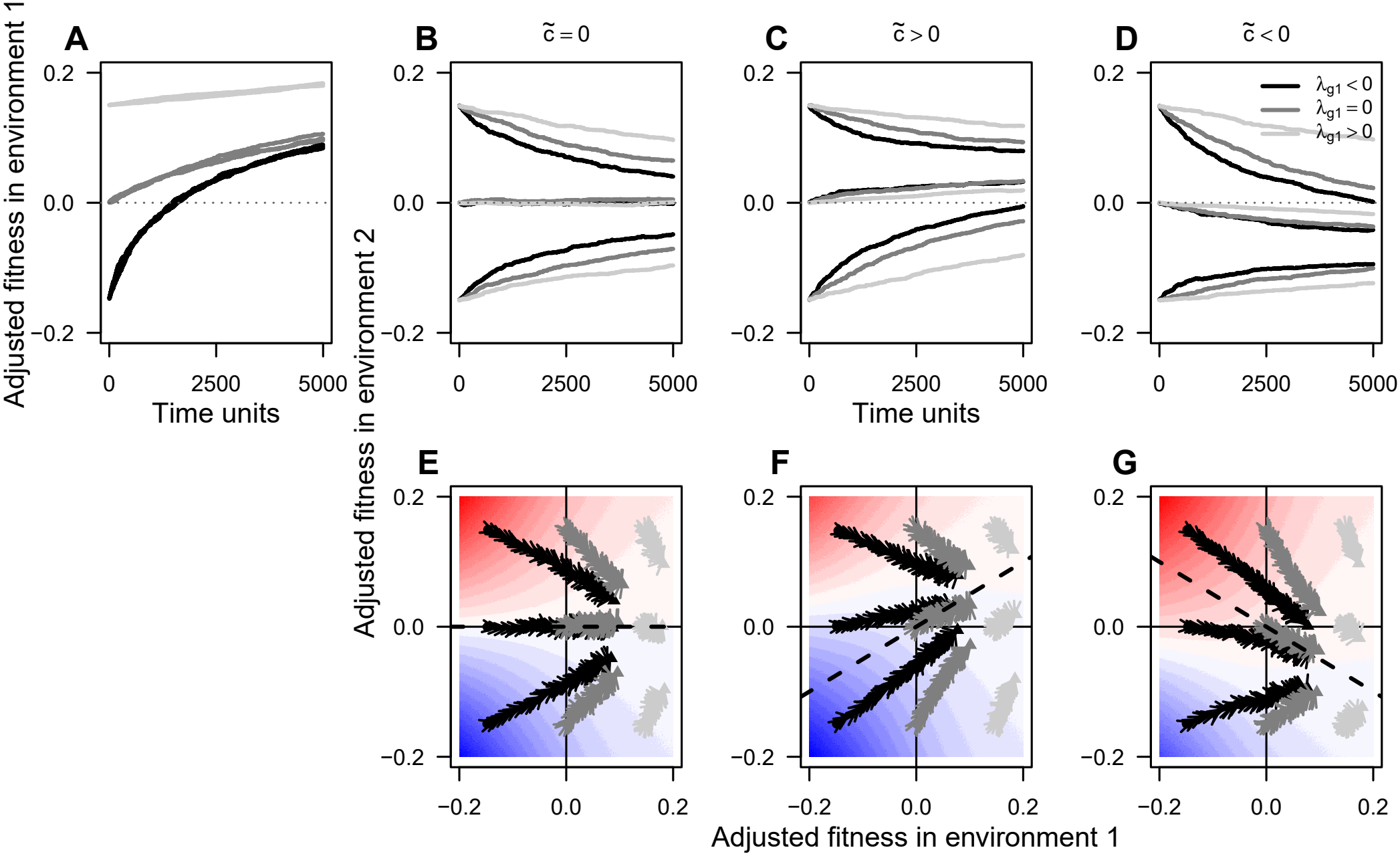
Predicted trajectories when global epistasis slopes are negatively correlated. Same parameters are used as main text Figure 4, except there is negative correlation between global epistasis slopes *ρ*_sl_ = −1. For no pleiotropic covariance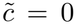, as predicted by our theory, the trajectories converge to the pivot fitness. For 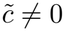 cases, the dashed line is no longer the attractor, nevertheless, the qualitative dynamics are similar to the same slope case.

**Figure S14.**
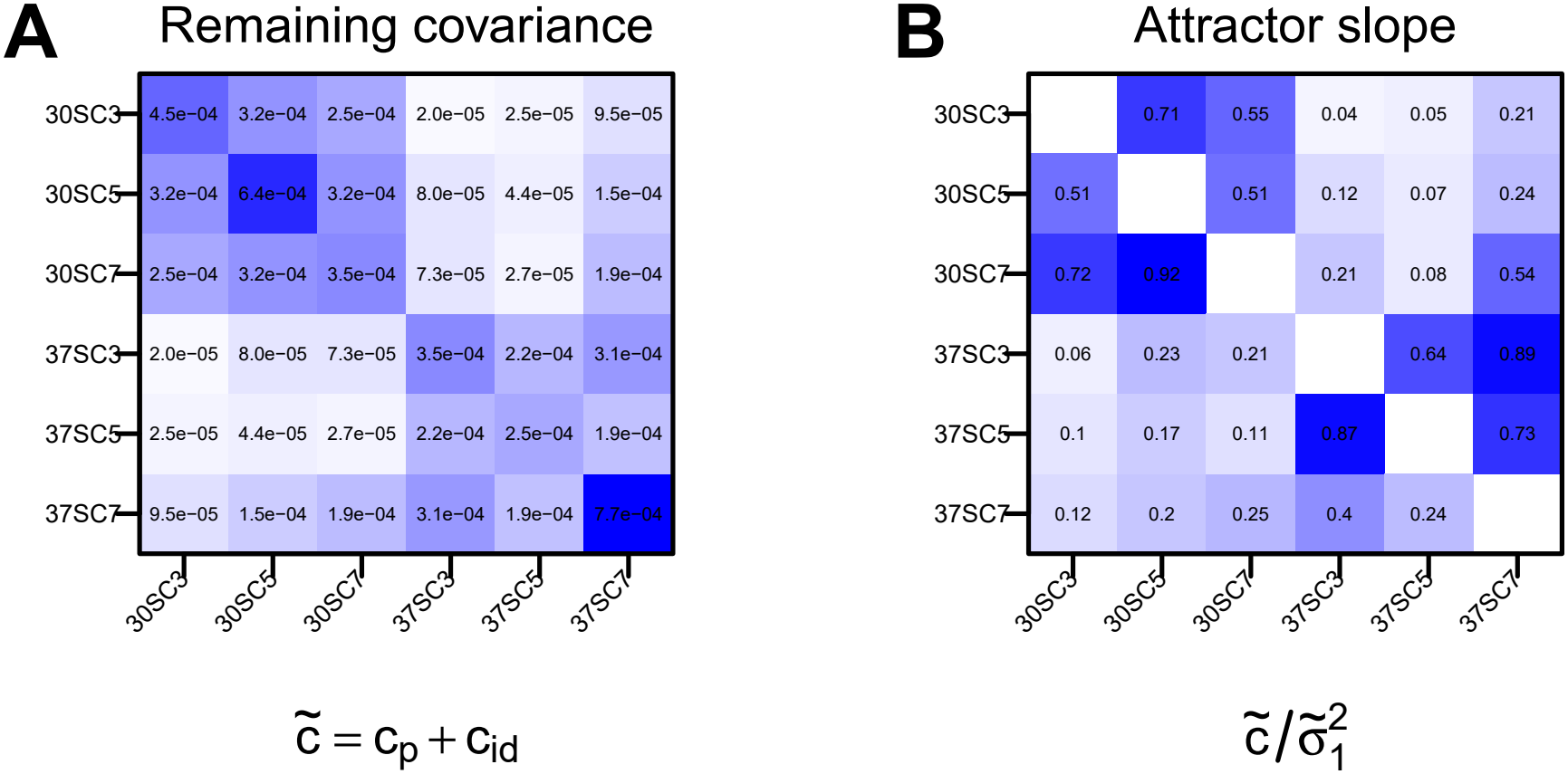
Theoretically predicted attractor slopes under the same slopes model for all ordered pairs of environments in the Ardell24 dataset. A: pleiotropic covariance 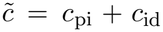 between environments. B: Predicted attractor slope 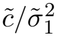. Rows indicate home environments and columns indicate non-home environments.

**Figure S15.**
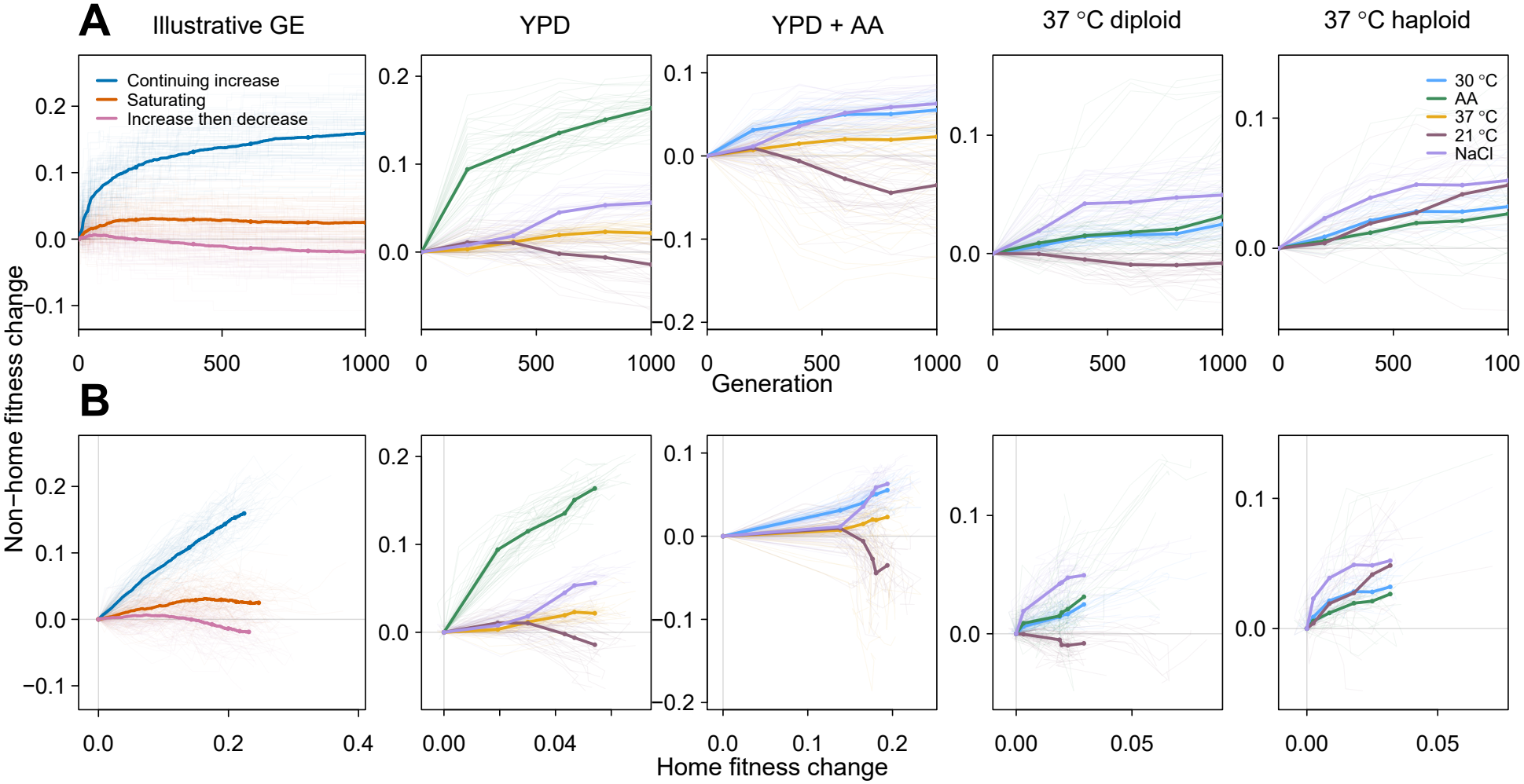
Our model captures qualitative dynamics of Bakerlee et al. (2021). A: Change in non-home fitness over time and B: Change in non-home fitness vs. change in home fitness. The first column shows simulations for the global epistasis model for three different initial non-home fitness and pleiotropic covariance 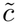 regimes. The remaining columns replot data from Bakerlee et al. (2021) for populations adapting in YPD at 30^*°*^C, YPD with 0.2% acetic acid (AA), YPD at 37^*°*^C (diploid), and YPD at 37^*°*^C (haploid). Faint lines represent individual replicate trajectories, while bold lines represent simulated means. Fitness changes are measured relative to initial fitness. The experimentally observed trajectories exhibit diverse qualitative dynamics that are consistent with the range of pleiotropic outcomes permitted by our model. Model parameters: Population size *N* = 10^4^, mutation rate *U* = 10^*−*5^, and *M* = 1000 initially sampled mutations. Mean global epistasis slopes *µ*_sl1_ = *µ*_sl2_ = −0.18, standard deviations *σ*_sl1_ = *σ*_sl2_ = 0.13, and slope correlation *ρ*_sl_ = 1. Mutations with positive slopes in either environment were excluded. Pivot noise standard deviations *σ*_pi1_ = *σ*_pi2_ = 0.015, with pleiotropic covariance 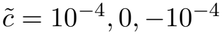 for the three regimes, respectively. Idiosyncratic epistasis was set to zero. Initial adjusted home fitness was 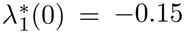 in all regimes; initial non-home fitness was defined as 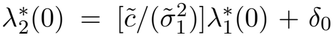, with *δ*_0_ = −0.08, −0.05, −0.10, respectively to adjust distance from theoretical attractor. Simulations were run for 1,000 time units with 100 replicate trajectories per regime.

## S2 Supplementary tables

**Table S1.** Same slope vs. different slopes model LRT.

| Model | df | logLik | AIC | BIC |
|---|---|---|---|---|
| Same slope | 46 | 78759.28 | -157426.55 | -157047.86 |
| Different slopes | 71 | 79104.02 | -158066.03 | -157481.52 |

**Table S2.**
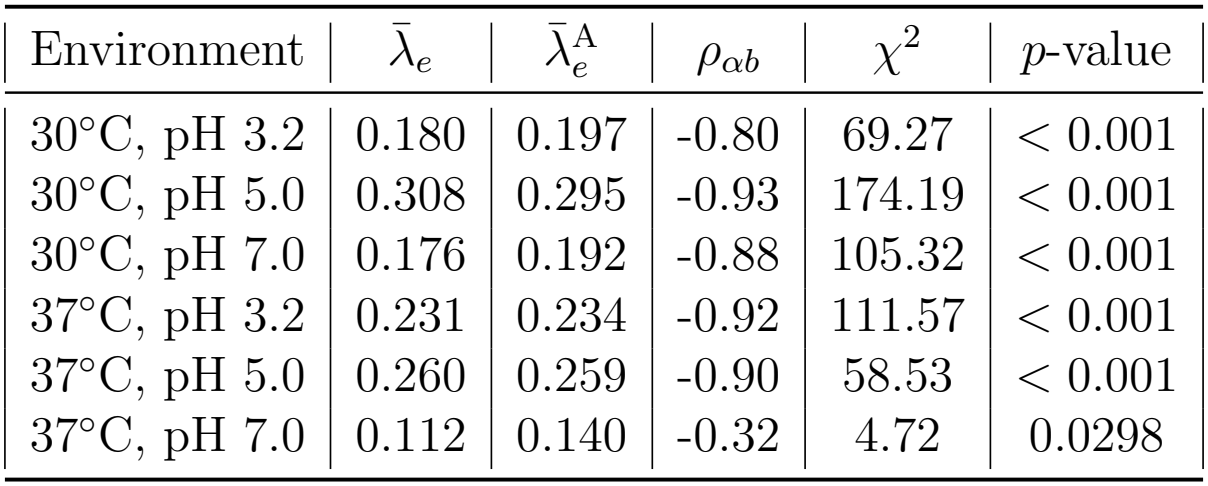
Estimated pivot growth rates and intercept-slope correlations. Pivot growth rates (in *h*^*−*1^) estimated in present study 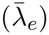 and Ardell et al. (2024) 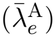, slope-intercept correlations within each environment (*ρ*_*αb*_). Likelihood-ratio tests compare models with and without slope-intercept covariance (*df* = 1).

| Environment | λ̄<sub>e</sub> | λ̄<sub>e</sub><sup>A</sup> | ρ<sub>αb</sub> | χ<sup>2</sup> | *p*-value |
|---|---|---|---|---|---|
| 30°C, pH 3.2 | 0.180 | 0.197 | -0.80 | 69.27 | < 0.001 |
| 30°C, pH 5.0 | 0.308 | 0.295 | -0.93 | 174.19 | < 0.001 |
| 30°C, pH 7.0 | 0.176 | 0.192 | -0.88 | 105.32 | < 0.001 |
| 37°C, pH 3.2 | 0.231 | 0.234 | -0.92 | 111.57 | < 0.001 |
| 37°C, pH 5.0 | 0.260 | 0.259 | -0.90 | 58.53 | < 0.001 |
| 37°C, pH 7.0 | 0.112 | 0.140 | -0.32 | 4.72 | 0.0298 |

**Table S3.**
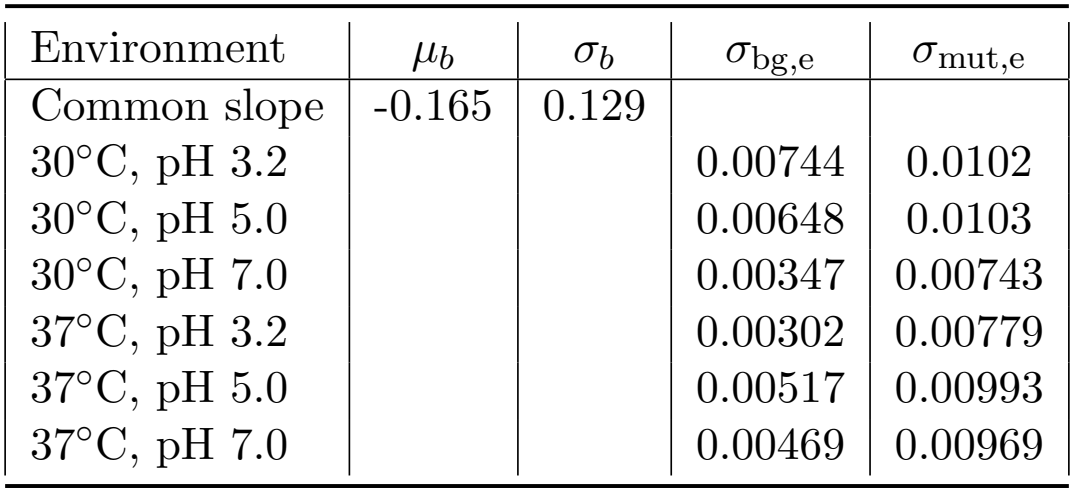
Parameter estimates for the same-slope global epistasis model. Fixed global epistasis slope (*µ*_*b*_), random slope s.d. (*σ*_*b*_), background growth rate error s.d. (*σ*_bg,e_), mutant growth rate error s.d. (*σ*_mut,e_). The fixed slope is significantly different from zero (Wald *z*-tests, *p <* 0.001). Remaining parameters are found in Figure S9.

| Environment | μ<sub>*b*</sub> | σ<sub>*b*</sub> | σ<sub>bg,e</sub> | σ<sub>mut,e</sub> |
|---|---|---|---|---|
| Common slope | -0.165 | 0.129 |  |  |
| 30°C, pH 3.2 |  |  | 0.00744 | 0.0102 |
| 30°C, pH 5.0 |  |  | 0.00648 | 0.0103 |
| 30°C, pH 7.0 |  |  | 0.00347 | 0.00743 |
| 37°C, pH 3.2 |  |  | 0.00302 | 0.00779 |
| 37°C, pH 5.0 |  |  | 0.00517 | 0.00993 |
| 37°C, pH 7.0 |  |  | 0.00469 | 0.00969 |

**Table S4.**
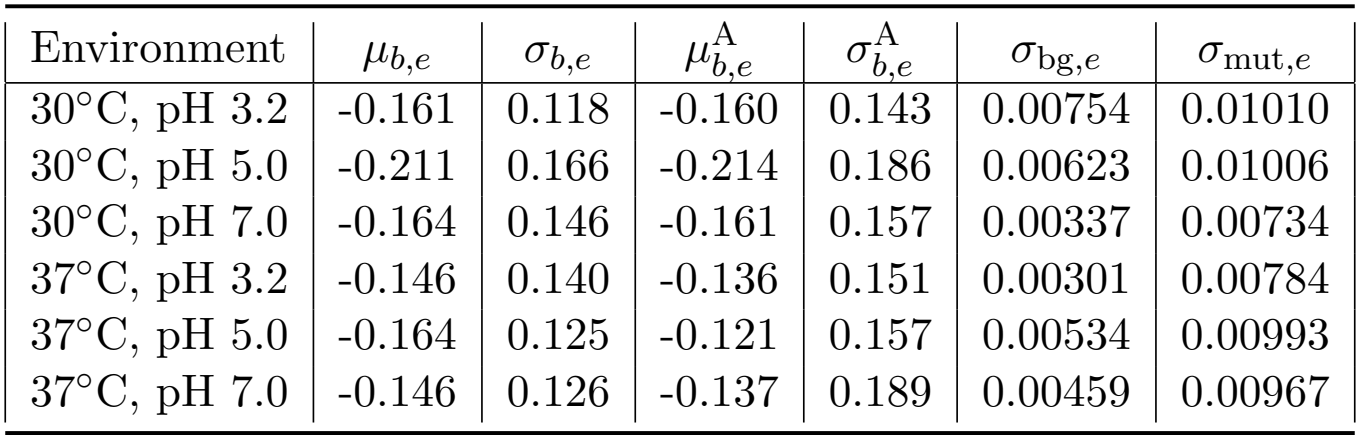
Parameter estimates for the different-slope global epistasis model. Environment-specific fixed global epistasis slopes (*µ*_*b,e*_), random slope s.d. (*σ*_*b,e*_), background growth rate measurement-error s.d. (*σ*_bg,*e*_), and mutant growth rate measurement-error s.d. (*σ*_mut,*e*_). The columns 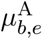 and 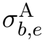 give the corresponding mean and s.d. of mutation-specific slopes reported by Ardell et al. (2024). All fixed slopes are significantly different from zero (Wald *z*-tests, *p <* 0.001). Remaining parameters are reported in SI Figure S6.

| Environment | *μ*<sub>*b*,*e*</sub> | *σ*<sub>*b*,*e*</sub> | $\mu_{b,e}^{\mathrm{A}}$ | $\sigma_{b,e}^{\mathrm{A}}$ | *σ*<sub>bg,*e*</sub> | *σ*<sub>mut,*e*</sub> |
|---|---|---|---|---|---|---|
| 30°C, pH 3.2 | -0.161 | 0.118 | -0.160 | 0.143 | 0.00754 | 0.01010 |
| 30°C, pH 5.0 | -0.211 | 0.166 | -0.214 | 0.186 | 0.00623 | 0.01006 |
| 30°C, pH 7.0 | -0.164 | 0.146 | -0.161 | 0.157 | 0.00337 | 0.00734 |
| 37°C, pH 3.2 | -0.146 | 0.140 | -0.136 | 0.151 | 0.00301 | 0.00784 |
| 37°C, pH 5.0 | -0.164 | 0.125 | -0.121 | 0.157 | 0.00534 | 0.00993 |
| 37°C, pH 7.0 | -0.146 | 0.126 | -0.137 | 0.189 | 0.00459 | 0.00967 |

## Notes

### Competing Interest Statement

The authors have declared no competing interest.

